# LRRC58 defines an E3 ubiquitin ligase complex sensitive to cysteine abundance

**DOI:** 10.1101/2025.09.23.678073

**Authors:** Dylan E. Ramage, Lianne H.E. Wieske, Charlotte Crowe, Jens B. Christensen, Theo A. von Wilmowski, Drew W. Grant, Zoe Bannister, Mark A. Nakasone, Kevin Haubrich, Mark Dorward, Iva A. Tchasovnikarova, Michael P. Weekes, N. Sumru Bayin, Alessio Ciulli, Richard T. Timms

**Affiliations:** Cambridge Institute for Therapeutic Immunology and Infectious Disease, Department of Medicine, University of Cambridge, Puddicombe Way, Cambridge, UK; Centre for Targeted Protein Degradation, School of Life Sciences, University of Dundee, Dundee, UK; The Gurdon Institute, University of Cambridge, Cambridge, UK; Department of Physiology, Development and Neuroscience, University of Cambridge, Cambridge, UK; Cambridge Institute for Medical Research, Department of Medicine, University of Cambridge, Cambridge, UK; Department of Biochemistry, University of Cambridge, Cambridge, UK

## Abstract

Adaptation to fluctuating nutrient supply is essential for organismal survival, but how cells sense and respond to changes in the abundance of many critical nutrients remains undefined. One such example is the amino acid cysteine, whose reactive thiol group is exploited for diverse cellular functions. Here, by characterizing the machinery required for the conditional degradation of cysteine dioxygenase type I (CDO1), the critical enzyme responsible for cysteine catabolism, we identify a Cul2 E3 ligase complex containing the uncharacterized substrate adaptor LRRC58 that is sensitive to cysteine abundance. When cysteine is replete, LRRC58 activity is restrained through auto-ubiquitination and proteasomal degradation; upon cysteine deprivation, LRRC58 is stabilized to permit CDO1 degradation. Through saturation mutagenesis stability profiling, we systematically validate a structural model of the CDO1-LRRC58 interaction and identify residues at the LRRC58 C-terminus required for cysteine-dependent instability. The LRRC58-mediated degradation of CDO1 is essential to prevent ferroptotic cell death under conditions of cysteine scarcity, and mutations in CDO1 which cause neurodevelopmental defects in humans encode dominant-active proteins refractory to LRRC58 recognition. Altogether, these data reveal the CDO1-LRRC58 axis as a critical regulator of cysteine homeostasis that safeguards neural development.

## INTRODUCTION

Simple organic compounds such as glucose, lipids and amino acids are an essential source of energy and the fundamental constituents of cellular biomass, and hence the ability to sense and respond to changes in nutrient availability is a prerequisite for life^1^. Homeostasis relies on a balance of nutrient uptake and catabolism as per energetic needs, and cells must therefore be able to both monitor nutrient levels and respond appropriately. However, despite the centrality of nutrient-sensing pathways throughout all kingdoms of life and their importance in the context of human disease, the underlying molecular mechanisms remain incompletely defined^2^.

A variety of signaling pathways are known to regulate cellular behavior in response to fluctuation in amino acid abundance. The kinase activity of GCN2 is activated upon binding to uncharged tRNA molecules that serve as a surrogate marker of amino acid insufficiency, resulting in the activation of the integrated stress response and upregulation of the transcription factor ATF4^3,4^. Most attention, however, has focused on dedicated sensors of specific amino acids which converge on the kinase activity of mechanistic target of rapamycin complex 1 (mTORC1), considered a master regulator of cell growth^5^. For example, the interactions between leucine and Sestrin2^6,7^, arginine and CASTOR^8,9^ and S-adenosylmethionine and SAMTOR^10,11^ regulate the behavior of the GATOR complexes which act as upstream regulators of mTORC1^12^. However, mTOR-independent mechanisms through which the abundance of specific amino acids regulates protein activity have also recently been uncovered; for example, the subcellular localization of HDAC6 is regulated by valine^13^ and the interaction between BAG2 and SAMD4B is regulated by arginine^14^. However, whether dedicated sensors specific for other essential amino acids also exist remains unknown.

We reasoned that conditional protein degradation via the ubiquitin-proteasome system (UPS) is another mechanism through which the cell could respond to altered amino acid availability. Ubiquitin is a 76-amino acid protein which is conjugated to target proteins through the sequential action of E1, E2 and E3 enzymes^15^, with the E3 ubiquitin ligases playing a key role in determining substrate specificity^16^. The potential consequences of protein ubiquitination are diverse^17^, but, as this modification can effect immediate change in the activity and/or stability of pre-existing proteins, the UPS is also ideally suited to enact rapid cellular responses upon fluctuations in nutrient supply. However, despite high-profile roles for E3 ligases in lipid-sensing pathways^18^, including the degradation of HMG-CoA reductase under sterol-replete conditions by RNF145 and gp78^19^ and the cholesterol-dependent ER-associated degradation (ERAD) of squalene monooxygenase by MARCHF6^20^, little is known about the role of the UPS in response to altered availability of amino acids.

Here, by adapting the Global Protein Stability (GPS) profiling approach^21,22^ to favor the identification of substrates subject to conditional degradation, we find that the stability of CDO1, the critical enzyme responsible for cysteine catabolism^23^, is correlated with cysteine abundance. A CRISPR/Cas9 genetic screen reveals a Cul2 complex comprising the substrate adaptor LRRC58 as the cognate E3 ligase responsible. Mechanistically, Cul2^LRRC58^ achieves conditional degradation of CDO1 through auto-regulation: in complete media, CDO1 remains stable as LRRC58 undergoes auto-ubiquitination and proteasomal degradation; conversely, as cysteine concentration falls, the auto-degradation of LRRC58 is inhibited allowing for CDO1 to be degraded. Saturation mutagenesis profiling of both CDO1 and LRRC58 validates a structural model of their interaction and identifies key residues at the C-terminus of LRRC58 that are essential for cysteine-dependent regulation. Finally, we show that the LRRC58-mediated regulation of CDO1 safeguards normal brain development: CDO1 mutations recently identified in children with severe neurological defects^24^ are dominant-active variants that prevent LRRC58-mediated regulation and promote cell death by ferroptosis. Altogether, these data demonstrate the importance of the CDO1-LRRC58 axis for cysteine homeostasis.

## RESULTS

### Comparative stability profiling reveals differential stability of CDO1

We set out to identify substrates of the UPS that are subject to conditional degradation. Reasoning that proteins exhibiting differential stability across cell types must, at some level, be subject to conditional regulation, we exploited the GPS expression screening platform^21,22^ to profile the stability of ∼14,000 human open reading frames (ORFs) across human cell lines. We began by performing stability profiling in Jurkat T cells **(Fig. S1A)**, comparing the results to a previous dataset^25^ generated in an analogous manner in HEK-293T cells **(Table S1)**. Although the vast majority of proteins exhibited concordant behavior between the two cell types, differential stability was observed for a handful of proteins **(Fig. 1A)**. The focus of this study was the strongest outlier, cysteine dioxygenase type 1 (CDO1), which was very stable in HEK-293T but unstable in Jurkat **(Fig. 1B-C)**.

**Figure 1.**
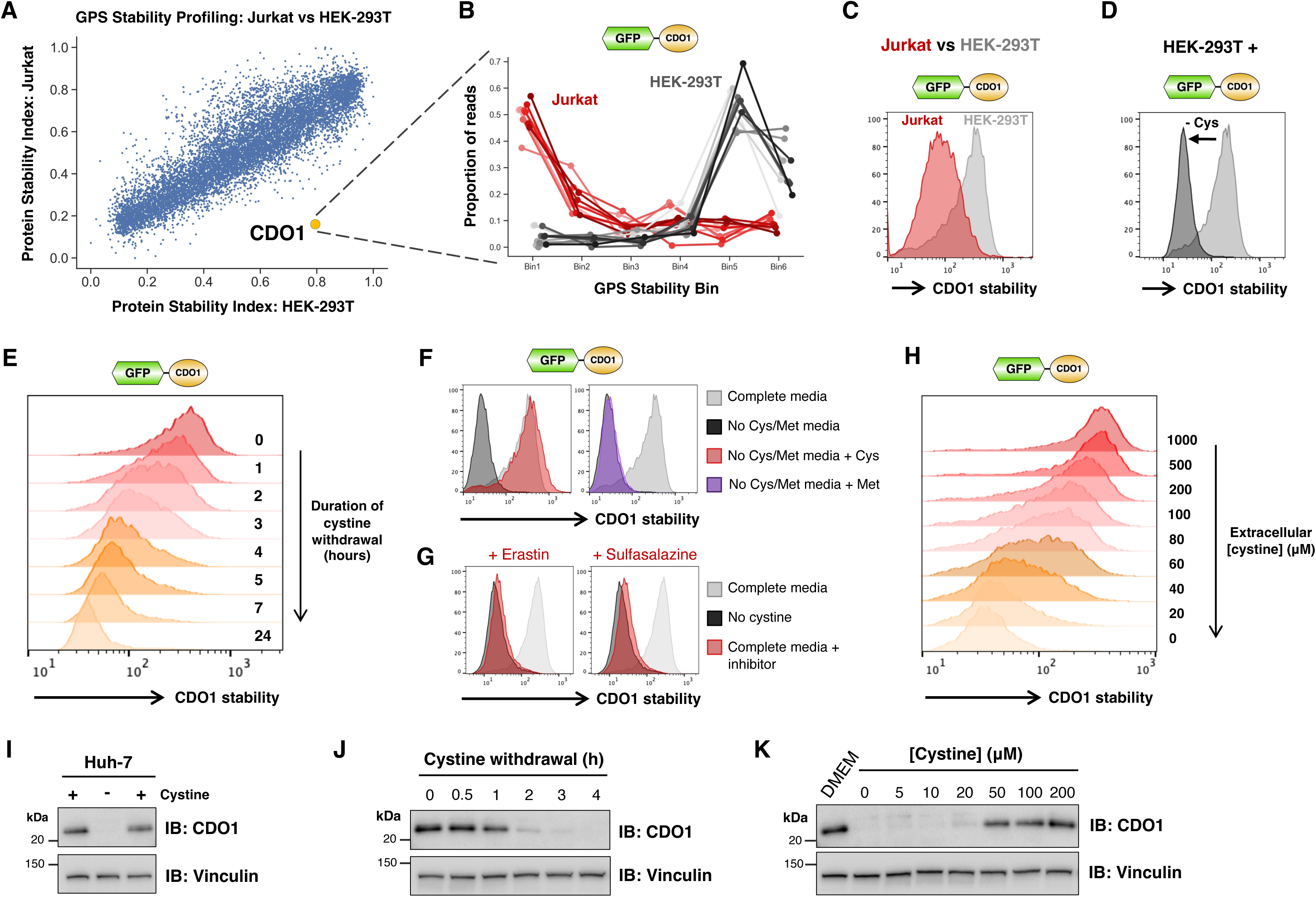
The stability of CDO1 is correlated with cysteine concentration. **(A-C)** CDO1 exhibits differential stability in Jurkat cells versus HEK-293T cells. **(A)** Stability profiling in Jurkat using the GPS ORFeome library. Each point represents the performance of an individual protein, with the stability (scaled between 0 = maximally unstable and 1 = maximally stable) in Jurkat (y-axis) compared to a previous dataset^25^ generated in HEK-293T cells (x-axis). **(B)** Stability profile of CDO1, which was represented by 10 barcoded replicates. **(C)** GPS-CDO1 was expressed in both HEK-293T and Jurkat cells and stability monitored by flow cytometry. **(D-G)** CDO1 is degraded upon cystine removal. CDO1 expressed in the context of the GPS system was expressed in HEK-293T cells and stability monitored by flow cytometry in the presence and absence of extracellular cystine and methionine **(D)**. A time course is shown in **(E)**. This effect was specifically due to withdrawal of cystine, as re-addition of cystine but not methionine prevented degradation **(F)**; furthermore, inhibition of cystine import via the xCT inhibitors erastin and sulfasalazine stimulated CDO1 degradation in complete media **(G)**. **(H)** The extent of CDO1 degradation is proportional to extracellular cystine concentration. HEK-293T cells expressing GPS-CDO1 were cultured overnight in cystine-free media reconstituted with the indicated concentrations of cystine and then analyzed by flow cytometry. **(I-K)** Endogenous CDO1 abundance is similarly regulated by cystine in Huh-7 cells, as assessed by immunoblot. Cystine withdrawal results in loss of CDO1 protein **(I)**; this effect occurs rapidly **(J)** and is correlated with extracellular cystine concentration **(K)**.

### CDO1 stability is regulated by cysteine abundance

CDO1 is the critical enzyme in cysteine catabolism, responsible for the oxidation of cysteine to cysteine sulfinic acid which ultimately leads to the production of taurine^23^ **(Fig. S1B)**. Cysteine is a particularly intriguing amino acid, as its reactive thiol group endows unique chemical properties to proteins^26^. Although sparingly used, cysteine residues play critical roles in both the structure and function of proteins, for example by allowing catalytic activity, disulfide bond formation and the coordination of metal ions^27^. Cysteine is also the rate-limiting substrate for the synthesis of glutathione, the key cellular antioxidant, and a precursor for essential sulfur-containing biomolecules including coenzyme A and hydrogen sulfide^28^.

Over two decades ago it was noted that CDO1 levels are modulated by cysteine abundance^29^, and thus we wondered whether relative cysteine levels might underlie the differential stability that we observed between HEK-293T cells and Jurkat cells. Indeed, upon withdrawal of exogenous cystine (the oxidized dimeric form of cysteine) and methionine from the culture media, GPS-CDO1 was rapidly degraded in HEK-293T cells **(Fig. 1D-E and Fig. S1C)**. CDO1 degradation was prevented upon restoration of extracellular cystine **(Fig. 1F)**, but, consistent with recent findings that many cancer cell lines lack the ability to synthesize cysteine *de novo* via the transsulfuration pathway^30^, re-addition of methionine had no effect **(Fig. 1F)**. Further supporting the notion that this effect was specific to depletion of cysteine, inhibition of the cystine transporter xCT with sulfasalazine^31^ or erastin^32^ also led to CDO1 degradation **(Fig. 1G)**, whereas mTOR inhibition^33^ or the addition of redox stressors did not **(Fig. S1D)**. Moreover, cystine supplementation alone was sufficient to abolish CDO1 degradation upon total amino acid withdrawal **(Fig. S1E)**. Furthermore, a catalytically-inactive CDO1 mutant (R60A) incapable of cysteine catabolism^34^ was degraded with slower kinetics than the wild-type protein **(Fig. S1F)** and, strikingly, the stability of the GPS-CDO1 fusion construct was proportional to the extracellular cystine concentration **(Fig. 1H)**. A similar mechanism appeared to operate in Jurkat, as cystine supplementation restored stability of the GPS-CDO1 fusion construct in Jurkat cells in a concentration-dependent manner **(Fig. S1G)**.

CDO1 is highly expressed in liver^35^, and, amongst a panel of commonly used cell lines, we were only able to detect CDO1 protein expression in the hepatocellular carcinoma cell lines Huh-7 and HepG2 **(Fig. S1H)**. Examining the fate of endogenous CDO1 in these two cell lines, we found that withdrawal of exogenous cystine led to a rapid decrease in CDO1 abundance **(Fig. 1I-J and Fig. S1I-J)**; moreover, CDO1 abundance was also correlated with extracellular cystine concentration **(Fig. 1K and Fig. S1K)**. Altogether, these data demonstrate that the stability of CDO1 is proportional to cysteine abundance, suggesting the existence of a cysteine-sensitive degradative pathway governing CDO1 degradation.

### Cul2^LRRC58^ and Cul5^LRRC58^ E3 ligase complexes degrade CDO1 in the absence of cysteine

We exploited our fluorescent GPS-CDO1 fusion construct to define the machinery responsible for CDO1 degradation. CDO1 degradation upon cystine withdrawal was blocked by the proteasome inhibitor Bortezomib and the E1 inhibitor TAK-243, indicating ubiquitin-dependent proteasomal degradation **(Fig. S2A)**. CDO1 degradation was also blocked by inhibition of Cullin-RING E3 ligases via treatment with MLN4924 **(Fig. S2A)**; furthermore, the expression of dominant-negative Cullin constructs suggested the involvement of a Cul2 or Cul5 E3 ligase complex **(Fig. S2B)**. Concordantly, a CRISPR screen using a UPS-focussed sgRNA library **(Fig. S2C)** identified a Cul2 E3 ligase complex containing the uncharacterized substrate adaptor LRRC58 as the cognate E3 responsible for CDO1 degradation **(Fig. 2A and Table S2)**. The domain architecture of LRRC58 reveals a series of leucine-rich repeats (LRRs), followed by BC-box and Cul-box motifs^36^ responsible for interactions with ElonginB/C and Cul2 respectively and a short, ordered C-terminal region **(Fig. 2B)**. CDO1 and LRRC58 have been identified as interacting partners through large-scale interactomics^37^, and AlphaFold 3^38^ predicts with high confidence (ipTM = 0.84) an interaction between CDO1 and LRRC58, with the putative interface mostly involving the LRRs **(Fig. 2C-D)**.

**Figure 2.**
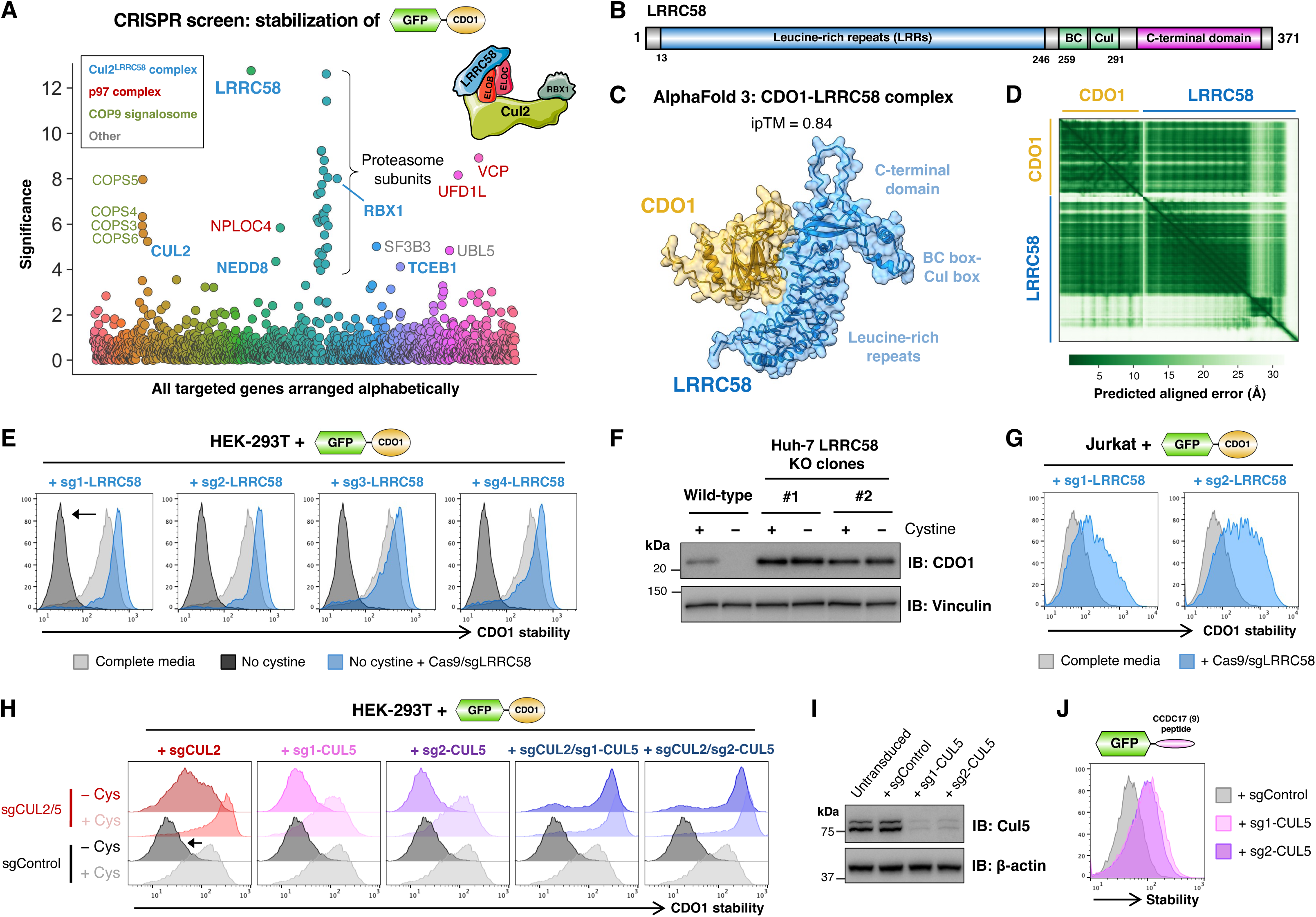
CDO1 is targeted for proteasomal degradation by the Cul2^LRRC58^ E3 ligase complex upon cysteine scarcity. **(A)** A CRISPR screen identifies a Cul2 complex containing the substrate adaptor LRRC58 as the E3 ligase responsible for CDO1 degradation. HEK-293T cells expressing GPS-CDO1 were transduced with a CRISPR sgRNA library targeting ubiquitin system components, and mutant GFP^bright^ cells unable to degrade CDO1 upon cystine withdrawal were isolated by FACS. “Significance” on the y-axis represents the negative log of the “pos|score” metric reported by the MAGeCK algorithm^52^. **(B)** Schematic representation of the domain architecture of LRRC58. (BC, BC-box; Cul, Cul-box) **(C-D)** AlphaFold 3 predicts an interaction between CDO1 and LRRC58 at high confidence. The structural prediction is shown in **(C)** with the predicted aligned error scores represented as a heatmap in **(D)**. **(E-G)** LRRC58 is responsible for CDO1 degradation. Individual CRISPR/Cas9-mediated gene disruption experiments validating the requirement for LRRC58 for CDO1 degradation upon cystine removal in HEK-293T cells as assessed by flow cytometry **(E)** and in HepG2 cells as assessed by immunoblot **(F)**. CRISPR/Cas9-mediated ablation of LRRC58 similarly stabilized CDO1 in Jurkat cells **(G)**. **(H-J)** Partial redundancy between Cul2 and Cul5 complexes for the LRRC58-mediated degradation of CDO1. **(H)** Individual CRISPR/Cas9-mediated disruption of *CUL2*, but not *CUL5*, partially abrogated CDO1 degradation upon cystine withdrawal; combined ablation of *CUL2* and *CUL5* was required to block CDO1 degradation. The sgRNA against *CUL2* was validated previously^22^; efficient disruption of *CUL5* was validated by immunoblot **(I)** and functionally through the stabilization of GFP fused to the peptide substrate CCDC17(9) targeted by Cul5^ASB7^ ^39^ **(J)**.

We validated the requirement for LRRC58 through individual CRISPR/Cas9-mediated gene disruption experiments. In HEK-293T cells LRRC58 ablation stabilized the GFP-CDO1 fusion protein at steady-state and completely inhibited its degradation upon cystine withdrawal **(Fig. 2E)**, while in Huh-7 cells LRRC58 ablation prevented the loss of endogenous CDO1 protein upon cystine withdrawal as assessed by immunoblot **(Fig. 2F)**. The same degradative mechanism was operational in Jurkat, where CRISPR-mediated LRRC58 disruption also prevented the proteasomal degradation of GPS-CDO1 **(Fig. 2G and Fig. S2D)**. We were unable to validate efficient CRISPR-mediated ablation of LRRC58 by immunoblot owing to the lack of effective commercial antibodies, but we demonstrated the specificity of these effects by confirming that exogenous expression of LRRC58 restored CDO1 degradation in an LRRC58 knockout clone **(Fig. S2E-F)**. Thus, in the absence of cysteine, CDO1 is targeted for proteasomal degradation via LRRC58.

Large-scale interactomics data^37^ suggests that LRRC58 associates with Cul5, but, unexpectedly, an interaction with Cul2 has not been reported. In support of our CRISPR screen results, however, we found that while CRISPR/Cas9-mediated ablation of *CUL2* did partially abrogate CDO1 degradation upon cystine deprivation, individual ablation of *CUL5* had no effect **(Fig. 2H)**. This finding could not be explained by inefficient *CUL5* disruption: both sgRNAs substantially reduced Cul5 protein abundance **(Fig. 2I)** and stabilized a reporter substrate targeted by Cul5^ASB7^ ^39^ **(Fig. 2J)**. Moreover, simultaneous CRISPR/Cas9-mediated ablation of both *CUL2* and *CUL5* completed blocked CDO1 degradation following cystine withdrawal, suggesting partially redundant roles **(Fig. 2H)**. Thus, although Cul2 complexes may be dominant, CDO1 is subject to degradation by both Cul2^LRRC58^ and Cul5^LRRC58^ E3 ligase complexes in the absence of cysteine.

### Saturation mutagenesis stability profiling validates a structural model of the CDO1-LRRC58 interaction

To evaluate the structural model of the CDO1-LRRC58 complex in an unbiased manner, we performed a deep mutational scan to identify substitution mutations which prevent CDO1 degradation following cystine withdrawal. In the context of the lentiviral GPS vector, we created a library of CDO1 variants in which each amino acid was systematically mutated to all other possible residues (200 residues x 20 amino acids = 4,000 variants) **(Fig. S3A)**. Upon single copy expression of the mutant library in HEK-293T cells, we profiled the stability of each of the resulting variants following cystine withdrawal by FACS combined with Illumina sequencing **(Fig. S3A and Table S3)**. The resulting data are summarized as a set of heatmaps in **Figure 3A**, where each cell represents the performance of an individual substitution mutant compared to the wild-type protein: the darker the red color, the greater the degree of CDO1 stabilization conferred by the mutation. Strikingly, mapping the stability scores onto the predicted structure of the CDO1-LRRC58 complex revealed that all of the residues identified as most critical for CDO1 degradation by the genetic screen lay on the predicted interaction interface **(Fig. 3B)**.

**Figure 3.**
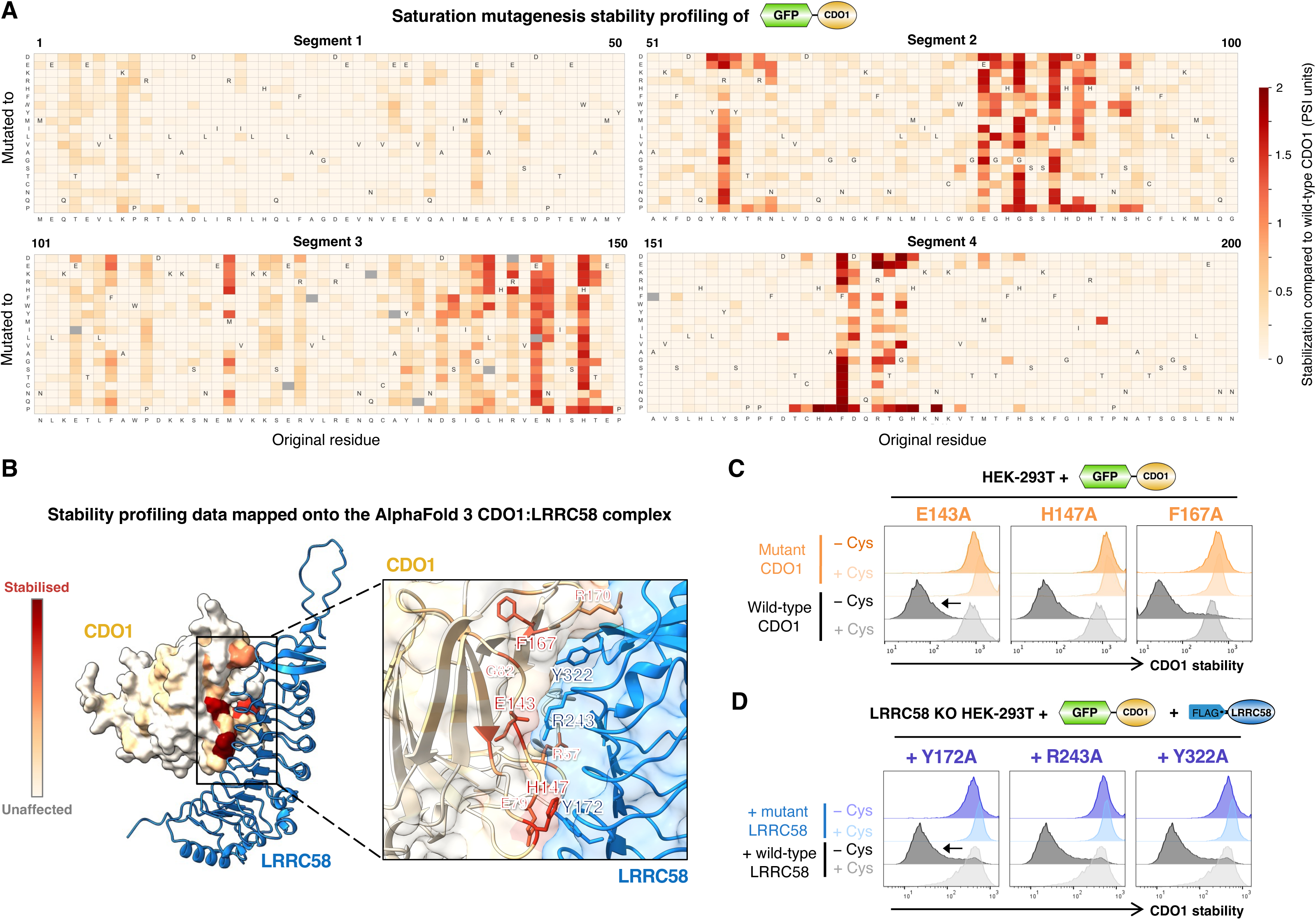
Saturation mutagenesis stability profiling of CDO1 validates a structural model of the CDO1-LRRC58 interaction. **(A)** Saturation mutagenesis stability profiling identifies substitution mutations which prevent CDO1 degradation upon cystine withdrawal. Each cell represents the stability of an individual substitution mutant; the darker the red color, the greater the degree of stabilization compared to the performance of wild-type CDO1. See also Fig. S3A. **(B)** Mutations which abrogate CDO1 degradation lie on the predicted interface with LRRC58. Mean stability scores for each residue of CDO1 from (A) were mapped onto the predicted structure of the CDO1-LRRC58 complex. **(C)** Individual validation of critical residues required for CDO1 degradation. The indicated CDO1 mutants were expressed in HEK-293T cells in the context of the GPS system, and stability monitored by flow cytometry in the presence and absence of cystine. **(D)** Assessing the requirement for the predicted interface residues in LRRC58. Wild-type CDO1 was expressed in LRRC58 KO HEK-293T cells in the context of the GPS system, and the ability of the indicated LRRC58 mutants to restore CDO1 degradation assessed by flow cytometry in the presence and absence of cystine.

We further validated these findings by examining the impact of a panel of individual point mutants. Mutation of CDO1 residues lying on the predicted interface with LRRC58 either abrogated or abolished CDO1 degradation upon cystine withdrawal **(Fig. 3C and Fig. S3B)**. To evaluate the impact of mutating the corresponding residues in LRRC58, we exploited our genetic complementation assay whereby the degradation of GPS-CDO1 in LRRC58 knockout cells could be restored upon exogenous expression of LRRC58 **(Fig. S2F)**. Concordantly, we found that LRRC58 mutants in which putative CDO1-interacting residues were replaced with alanine did not support CDO1 degradation upon cystine withdrawal **(Fig. 3D)**. These phenotypes were indeed due to a loss of the CDO1-LRRC58 interaction, as assessed through co-immunoprecipitation experiments **(Fig. S3C-D)**. Therefore, we concluded that the structural prediction represents a faithful model of the CDO1-LRRC58 interaction.

### The stability of LRRC58 is inversely correlated with cysteine abundance

The goal of our subsequent experiments was to understand how the degradation of CDO1 by Cul2^LRRC58^ is conditionally regulated by cysteine abundance. We first considered the possibility that CDO1 could act as a cysteine “sensor”; however, active site mutations predicted to abolish cysteine binding had no effect on CDO1 stability in our deep mutational scan **(Fig. 3)**, and, although the crystal structure of CDO1 bound to cysteine^40^ suggests a potential second cysteine binding site on the surface of CDO1, mutations targeting this region also did not affect CDO1 stability **(Fig. 3)**. We then considered whether cysteine abundance might regulate the interaction between CDO1 and LRRC58, but co-immunoprecipitation experiments suggested that GFP-CDO1 could bind FLAG-LRRC58 in both the presence and absence of cysteine (**Fig. S4A)**. However, this experiment did reveal that the abundance of exogenous LRRC58 was increased in the absence of cysteine (**Fig. S4A, left panel)**. Indeed, when examined in the context of the GPS system, we found that LRRC58 was highly unstable in complete media but was stabilized upon withdrawal of extracellular cystine **(Fig. 4A)**. This effect was mirrored upon inhibition of cystine import using sulfasalazine or erastin **(Fig. 4B)**, but not by mTOR inhibition or other redox stressors **(Fig. S4B)**. LRRC58 stabilization occurred rapidly following cystine withdrawal **(Fig. 4C)**, and this effect was not a consequence of the GFP-fusion partner, as an HA-tagged LRRC58 construct exhibited similar behavior when monitored by immunoblot **(Fig. 4D)**. Notably, when we examined the proteome globally using tandem mass tag (TMT) proteomics **(Table S4)**, LRRC58 emerged as the protein exhibiting the most substantial increase in abundance following cystine depletion **(Fig. 4E)**. Most strikingly, the stability of GPS-LRRC58 was inversely correlated with the extracellular cystine concentration **(Fig. 4F-G)**, as was the abundance of HA-tagged LRRC58 when assessed by immunoblot **(Fig. 4H)**. Altogether, we concluded that cysteine availability regulates CDO1 degradation at the level of LRRC58 stability; furthermore, as HEK-293T cells do not express CDO1, these data demonstrated that LRRC58 can respond independently to altered cysteine abundance.

**Figure 4.**
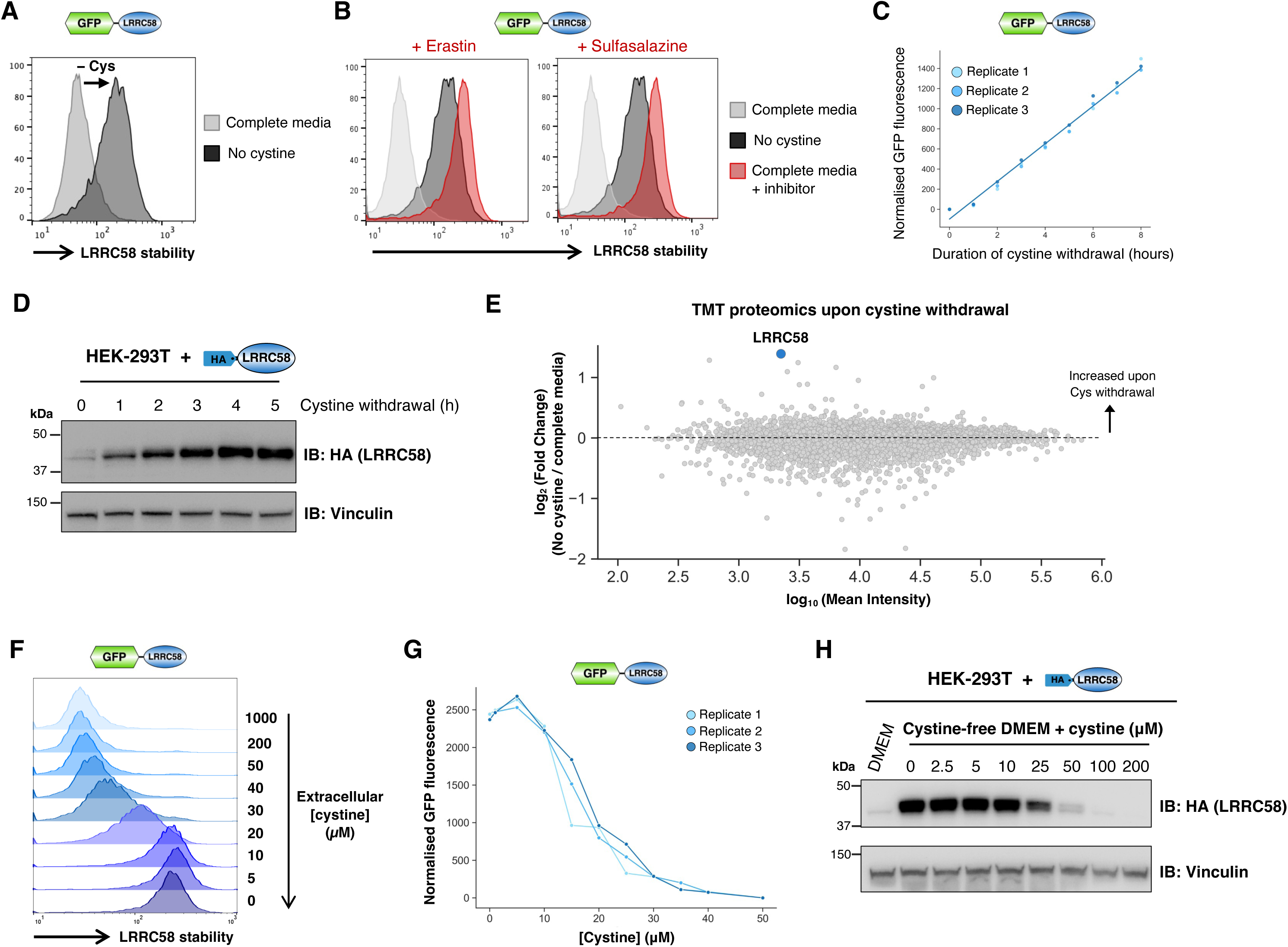
The stability of LRRC58 is inversely correlated with cysteine abundance. **(A-D)** LRRC58 is stabilized upon cystine withdrawal. **(A)** LRRC58 was expressed in HEK-293T cells in the context of the GPS system, and stability monitored by flow cytometry in the presence and absence of extracellular cystine. LRRC58 was also stabilized in complete media upon inhibition of cystine import **(B)**. A time course monitoring the stabilization of GPS-LRRC58 following cystine withdrawal by flow cytometry is shown in **(C)**; analogous data monitoring HA-LRRC58 following cystine withdrawal by immunoblot is shown in **(D)**. **(E)** TMT proteomics measuring global protein abundance 12 hours after cystine and methionine deprivation; LRRC58 exhibits the greatest relative increase in abundance. **(F-H)** LRRC58 stability is inversely correlated with cysteine abundance. **(F)** HEK-293T cells expressing GPS-LRRC58 were cultured overnight in cystine-free media reconstituted with the indicated concentrations of cystine and then analyzed by flow cytometry; quantitation of GFP fluorescence is shown in **(G)**. The abundance of HA-LRRC58 was also inversely correlated with extracellular cystine concentration, as monitored by immunoblot **(H)**.

### Auto-ubiquitination underlies LRRC58 instability in the presence of cysteine

We extended these observations by examining the mechanistic basis for LRRC58 instability when cysteine is replete. GFP-tagged LRRC58 expressed in the context of the GPS system was stabilized upon proteasome and E1 inhibition **(Fig. S4C)**, indicating ubiquitin-mediated proteasomal degradation. Furthermore, GPS-LRRC58 was also stabilized by Cullin-RING ligase inhibition **(Fig. S4C)**, expression of dominant-negative Cul2 **(Fig. S4D)** and CRISPR/Cas9-mediated disruption of *CUL2* **(Fig. S4E)**, indicating the involvement of a Cul2 E3 ligase complex. However, a UPS-focused CRISPR screen designed to identify genes required for LRRC58 instability in complete media revealed no significant enrichment of any other candidate Cul2 substrate adaptors aside from LRRC58 itself **(Fig. S4F and Table S5)**. Altogether, we considered that the most plausible hypothesis was that LRRC58 regulates its own stability via auto-ubiquitination. Supporting this notion, we found that deletion of either the BC-box or the Cul-box, motifs responsible for ElonginB/C and Cul2 binding respectively, rendered LRRC58 stable in the presence of cysteine **(Fig. 5A)**. This effect was recapitulated with individual point mutants in these motifs **(Fig. 5B)**, which we validated prevented association with Cul2 **(Fig. 5C)**.

**Figure 5.**
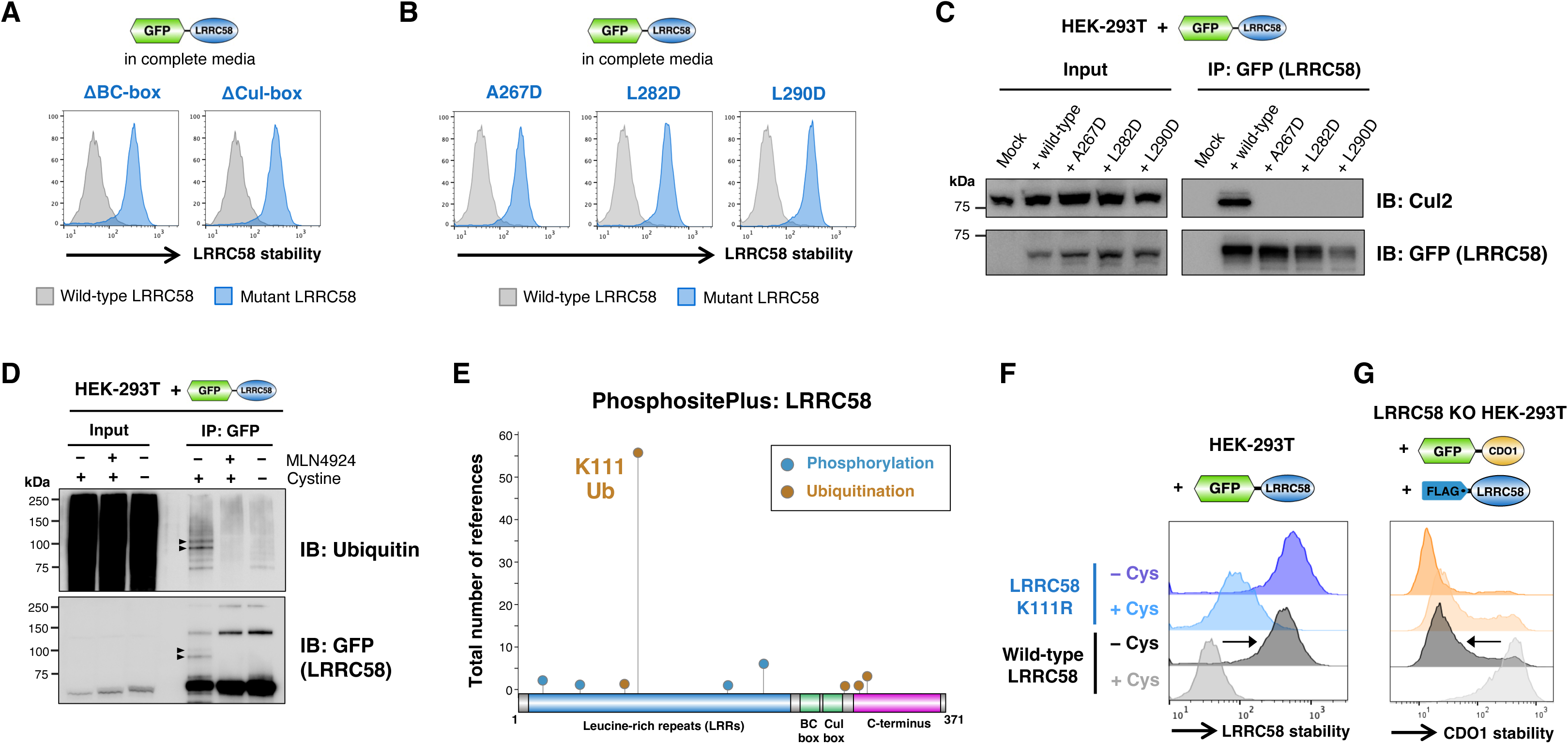
Auto-ubiquitination underlies the instability of LRRC58 in the presence of cysteine. **(A-C)** LRRC58 mutants unable to associate with the Cul2 complex remain stable in complete media. **(A)** LRRC58 deletion mutants lacking either the BC-box or Cul-box motifs did not exhibit instability in complete media when assessed in the context of the GPS system by flow cytometry. Point mutations targeting individual residues within the BC-box and Cul-box motifs exerted the same stabilizing effect **(B)** and could not associate with Cul2 as assessed via co-immunoprecipitation **(C)**. **(D)** LRRC58 is ubiquitinated in the presence of cysteine. GFP-tagged LRRC58 was immunoprecipitated from HEK-293T cells and ubiquitination assessed by immunoblot using the VU-1 antibody. Arrows indicate ubiquitinated species apparent in complete media but not following Cullin-RING ligase inhibition with MLN4924 or removal of cystine. **(E-G)** Ubiquitination at K111 restrains LRRC58 activity. **(E)** PhosphositePlus data revealing LRRC58 ubiquitination at K111 has been frequently observed in mass spectrometry experiments. An LRRC58 mutant unable to be ubiquitinated at this site (K111R) was partially stabilized in complete media **(F)**; this resulted in hyperactivity, as, unlike wild-type LRRC58, the K111R mutant degraded CDO1 efficiently in complete media **(G)**.

Biochemically, we detected ubiquitination of exogenous LRRC58 in cells grown in complete media, but this effect was abrogated upon Cullin-RING ligase inhibition with MLN4924 or withdrawal of exogenous cystine **(Fig. 5D)**. Intriguingly, many proteomic studies have reported ubiquitination of LRRC58 at lysine-111 (K111) **(Fig. 5E)**, suggesting that this residue may be the primary ubiquitin acceptor site responsible for LRRC58 instability. Indeed, a LRRC58 mutant unable to be ubiquitinated at this site (K111R) was considerably more stable than wild-type LRRC58 in the presence of cysteine **(Fig. 5F)**. Strikingly, the K111R mutant was functionally hyperactive compared to wild-type LRRC58, efficiently promoting the degradation of CDO1 even in the presence of cysteine **(Fig. 5G)**. Altogether, these data demonstrate that it is auto-regulation of LRRC58 abundance in response to cysteine availability that imparts conditionality to the degradation of CDO1.

### Critical residues at the LRRC58 C-terminus confer cysteine-dependent regulation

The data described in the previous section suggests that cysteine sufficiency stimulates the auto-ubiquitination and proteasomal degradation of LRRC58. With the goal of systematically defining the residues of LRRC58 required for this effect, we performed two deep mutational scans. To simplify this task, we focussed our attention on the C-terminus of LRRC58. Given that the leucine-rich repeats mediate the interaction with CDO1 and the BC-box and Cul-box motifs are responsible for the interaction with the Cullin complex, we reasoned that the C-terminal domain of LRRC58 was by far the most likely part of the protein to confer cysteine-dependent regulation; moreover, we found that even small deletions of the LRRC58 C-terminus abolished instability in complete media **(Fig. S4G)**.

We designed a pool of single amino acid variants in which each residue of LRRC58 from glutamine-156 (Q156) through to the C-terminal residue glycine-371 (G371) was mutated to all other possible residues (216 residues x 20 amino acids = 4,320 variants), and expressed this mutant library in two settings **(Fig. S5A)**: (1) in the context of the GPS expression vector, with the goal of identifying point mutations which stabilize LRRC58 in the presence of cysteine **(Table S6)**, and (2) in the complementation of LRRC58 KO cells expressing GPS-CDO1, to define residues which hyperactivate the degradation of CDO1 in the presence of cysteine **(Table S7)**. A global summary of the resulting data is presented as heatmaps in **Figures 6A-B** and **Figures S5B-C**, where red cells indicate LRRC58 mutants which are stabilized in the presence of cysteine, and blue cells indicate LRRC58 mutants which are hyperactive in the presence of cysteine.

**Figure 6.**
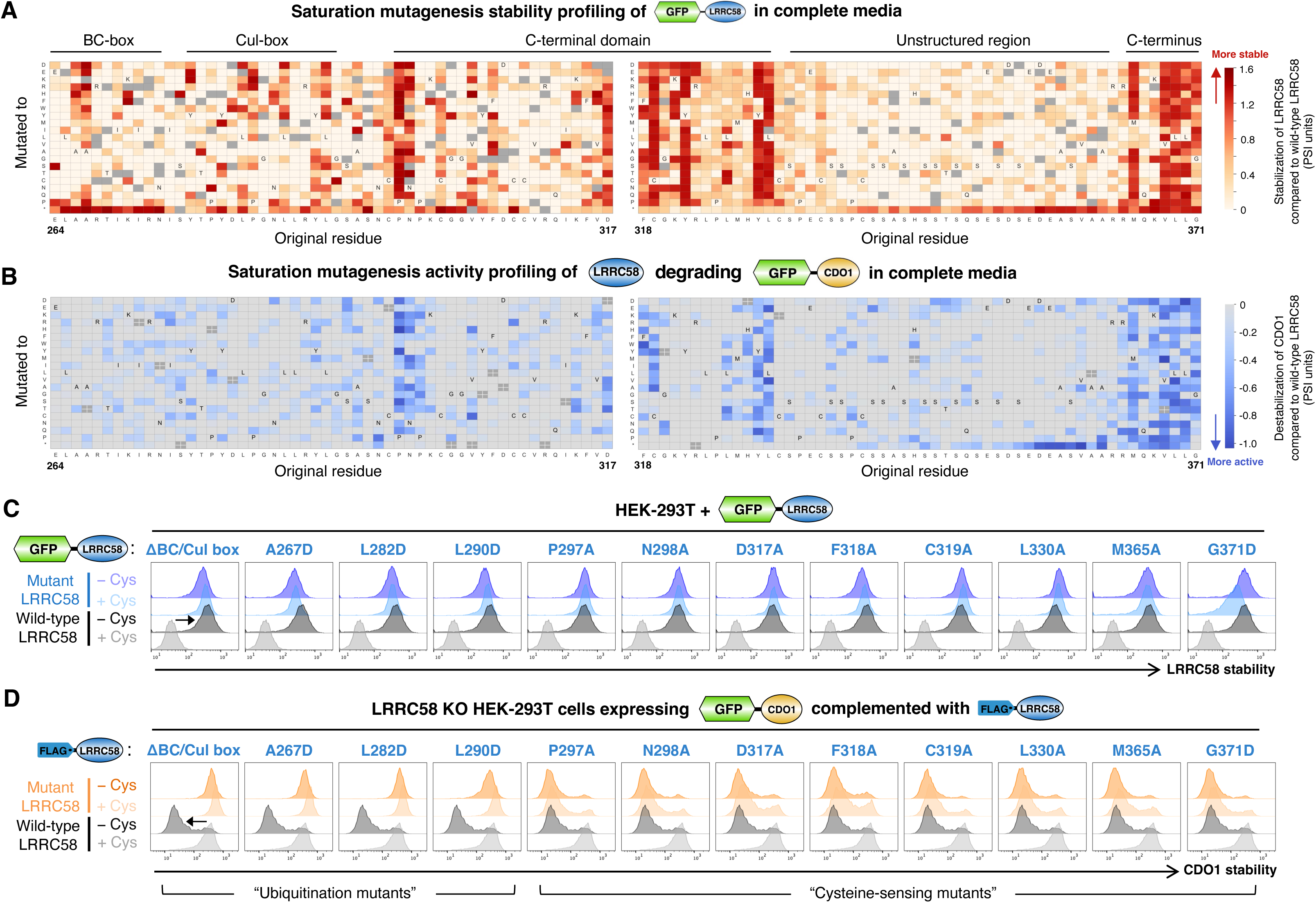
Twin deep mutational scans identify the critical residues of LRRC58 responsible for cysteine sensitivity. **(A)** Saturation mutagenesis stability profiling of LRRC58 in complete media. Each cell represents the performance of an individual substitution mutant; the darker the red color, the greater the degree of stabilization compared to wild-type LRRC58. Dark gray cells represent mutants for which insufficient data was available for analysis. See also Fig. S5A-B. **(B)** Saturation mutagenesis activity profiling of LRRC58, in which the ability of LRRC58 mutants to degrade CDO1 was assessed in complete media. Each cell represents the performance of an individual substitution mutant; the darker the blue color, the more hyperactive the mutant compared to wild-type LRRC58. See also Fig. S5A and S5C. **(C-D)** Individual validation experiments. **(C)** All of the mutants examined stabilized LRRC58 in complete media. **(D)** Two functional classes of mutant were apparent when assessing their ability to degrade CDO1: “ubiquitination mutants” (left) were unable to degrade CDO1 irrespective of cysteine abundance, while “cysteine-sensing mutants” (right) exhibited hyperactivity, efficiently degrading CDO1 in complete media.

The stability profiling screen revealed multiple insights into the control of LRRC58 stability. First, that the folded structure of LRRC58 is critical for instability in the presence of cysteine. In particular, substitutions replacing structural leucine, isoleucine and asparagine residues in the leucine-rich repeats with polar or charged amino acids conferred stabilization **(Fig. S5B)**, suggesting that the specific architecture of LRRC58 is necessary for auto-ubiquitination. Second, that the C-terminal domain of LRRC58 plays a central role in stability control, with a series of residues (P297, N298, D317, F318, C319, Y322, Y329 and L330) absolutely critical for cysteine-mediated instability **(Fig. 6A)**. Third, that the extreme C-terminus of LRRC58 also plays a vital role, with the data highlighting a specific terminal motif (-VLLG*) essential for cysteine-mediated instability **(Fig. 6A)**.

Comparing the stability profiling results with those from the LRRC58 activity scan **(Fig. 6B and Fig. S5C)** enabled us to classify the mutants which remain stable in the presence of cysteine into two functional classes: (1) mutants that are unable to mediate CDO1 degradation, either because of an inability to bind CDO1 (for example mutations in the LRRs predicted to disrupt the architecture of LRRC58) or because of defective ubiquitination (for example mutations which prevent the association between LRRC58 and the Cullin scaffold), and (2) “cysteine-sensing” mutants, which are competent to target CDO1 but fail to auto-regulate in the presence of cysteine, and hence efficiently degrade CDO1 irrespective of cysteine abundance.

We validated the screen results by individually examining the stability **(Fig. 6C)** and degradative activity **(Fig. 6D)** of a panel of representative LRRC58 mutants. All of the mutations tested led to stabilization of LRRC58 in the presence of cysteine **(Fig. 6C)**, while the two classes of mutant were apparent when assessing their ability to restore CDO1 degradation in LRRC58 KO cells **(Fig. 6D)**: the “ubiquitination mutants” A267D, L282D and L290D affecting the BC-box and Cul-box motifs were entirely non-functional and unable to degrade CDO1 even in the absence of cysteine, while the “cysteine-sensing” mutants P297A, N298A, D317A, F318A, C319A, L330, M365A and G371D exhibited hyperactivity, efficiently degrading CDO1 in complete media. Altogether, these data systematically define residues near the LRRC58 C-terminus that are required for cysteine-dependent regulation.

### The LRRC58-mediated degradation of CDO1 prevents ferroptosis upon cysteine scarcity

These data support a model whereby when cysteine is replete, CDO1 remains stable to promote cysteine breakdown, whereas when cysteine is scarce, CDO1 is rapidly degraded to preserve intracellular cysteine reserves. To test the importance of LRRC58 activity towards CDO1 under conditions of cysteine scarcity, we examined the fate of a panel of HEK-293T cell lines **(Fig. S6A-B)** upon cysteine withdrawal. The viability of wild-type HEK-293T cells, which do not express CDO1, was largely unaffected by overnight cystine withdrawal **(Fig. 7A, gray bars)**. The same was also true following exogenous expression of CDO1, but only in cells in which LRRC58 was active: both LRRC58 KO cells and LRRC58 KO cells reconstituted with an LRRC58 mutant (R243A) unable to bind CDO1 succumbed to overnight cystine withdrawal, whereas LRRC58 KO cells complemented with wild-type LRRC58 were protected **(Fig. 7A, gold bars)**. This effect was dependent on CDO1 activity, as cystine withdrawal was not toxic to cells expressing a catalytically-inactive CDO1 mutant (R60A) **(Fig. 7A, pink bars)**. Conversely, introduction of a CDO1 mutant (E143A) refractory to LRRC58 degradation was toxic upon cystine withdrawal regardless of LRRC58 status **(Fig. 7A, red bars)**.

**Figure 7.**
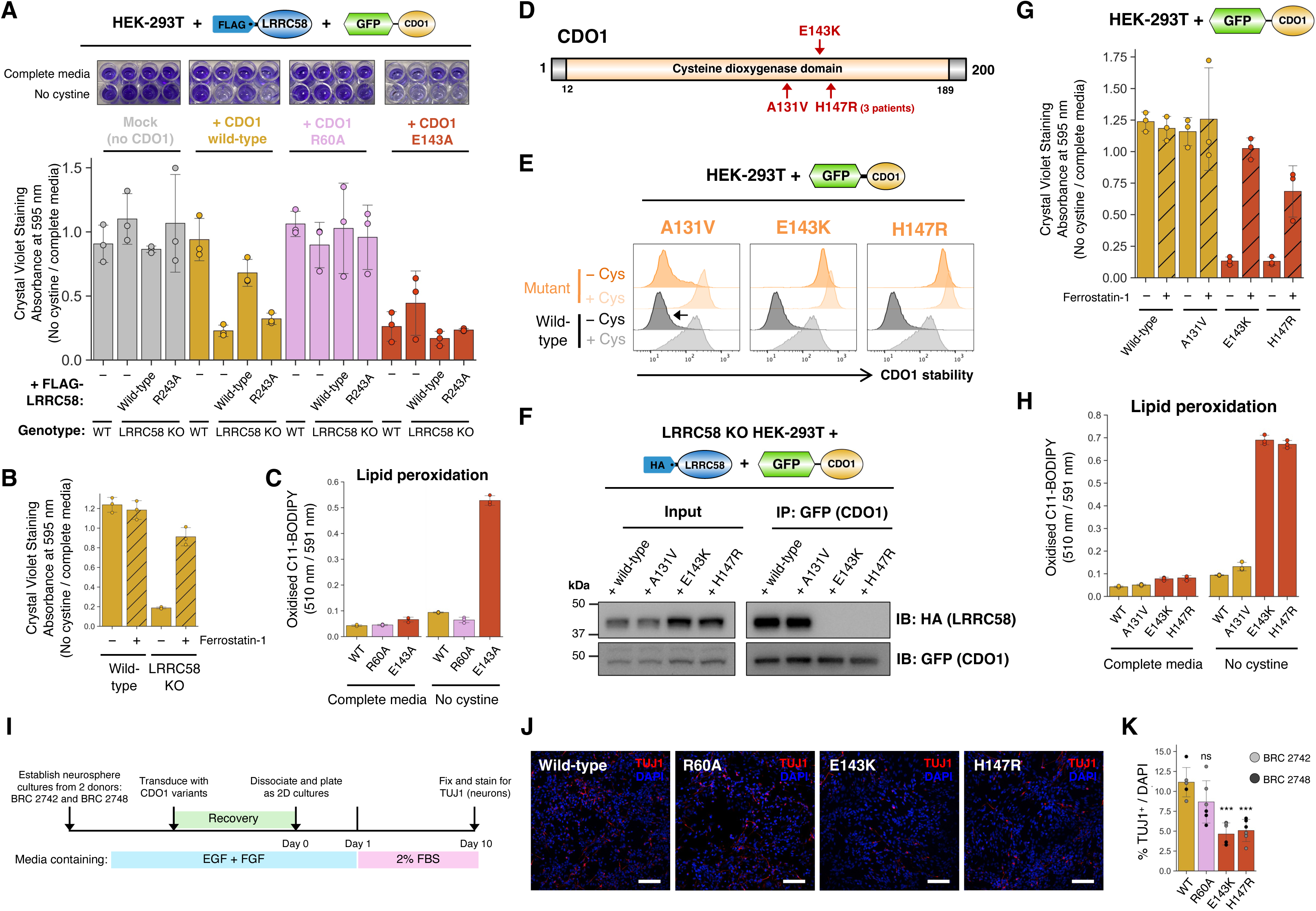
CDO1 mutations that impair brain development encode dominant-active proteins refractory to LRRC58-mediated degradation. **(A)** The LRRC58-mediated degradation of CDO1 is critical for cell viability under conditions of cysteine scarcity. A panel of HEK-293T cell lines were manipulated to modulate CDO1 and LRRC58 expression, and then cell viability following overnight incubation in cystine-free media was assessed by crystal violet staining. Cystine deprivation is toxic to cells expressing a catalytically-active CDO1 if unrestrained by LRRC58. See also Fig. S6A-B. **(B-C)** Loss of LRRC58 promotes ferroptotic cell death upon cystine starvation. **(B)** The death of LRRC58 KO cells expressing CDO1 following cystine withdrawal could be rescued by the ferroptosis inhibitor Ferrostatin-1 (hatched bars); moreover, C11-BODIPY staining **(C)** revealed a substantial increase in lipid peroxidation upon cystine deprivation in cells expressing the E143A mutant CDO1 which does not bind LRRC58. See also Fig. S6C. **(D)** Schematic representation of dominant mutations in CDO1 recently associated with severe neurodevelopmental abnormalities^24^. **(E-F)** CDO1 disease mutants encode dominant-active proteins that cannot be regulated by LRRC58. The E143K and H147R CDO1 disease mutants were not subject to degradation upon cystine withdrawal when examined in the context of the GPS system **(E)** and were unable to bind LRRC58 as assessed by co-immunoprecipitation **(F)**. **(G-H)** Expression of CDO1 disease mutants promotes ferroptotic cell death. **(G)** Crystal violet staining to assess the viability of HEK-293T cells after overnight cystine deprivation; the toxic effects resulting from expression of the E143K and H147R CDO1 mutants was abrogated by Ferrostatin-1 treatment. **(H)** C11-BODIPY staining to assess lipid peroxidation. Expression of the E143K and H147R CDO1 mutants resulted in a small increase in the proportion of oxidized C11-BODIPY in complete media, and a dramatic increase following cystine withdrawal. See also Fig. S6C. **(I-K)** Expression of CDO1 disease mutants impairs the differentiation of primary human neurosphere cultures. **(I)** Schematic representation of the experiment. Neurosphere cultures established from fetal brain tissue either nine (BRC 2742) or twelve (BRC 2748) weeks post conception were transduced with lentiviral vectors encoding CDO1 variants, induced to differentiate into mature neurons through the addition of 2% FBS, and then analyzed by immunofluorescence microscopy for TUJ1 expression. Representative images are shown in **(J)**, with the proportion of TUJI^+^ neurons generated quantified in **(K)**. (*** P < 0.001, Tukey’s HSD test compared to wild-type CDO1; ns, not significant. Scale bar, 100 µm)

Depletion of intracellular cysteine can lead to ferroptosis, a non-apoptotic form of cell death characterized by peroxidation of membrane phospholipids^41^. We found that the ferroptosis inhibitor Ferrostatin-1^42^ substantially ameliorated the toxicity associated with cystine depletion in LRRC58 KO cells **(Fig. 7B)**, and expression of the E143A CDO1 mutant resulted in a dramatic increase in lipid peroxidation prior to cell death as assessed using the C11-BODIPY dye^43^ **(Fig. 7C and Fig. S6C)**. Thus, these data suggest that the ability of LRRC58 to restrain cysteine catabolism via CDO1 degradation is necessary to prevent ferroptotic cell death under conditions of cysteine scarcity.

### Dominant-active CDO1 mutants unable to bind LRRC58 impede neural development

Dominant heterozygous missense mutations in *CDO1* were recently proposed as the underlying cause of severe neurodevelopmental abnormalities in five children^24^. Three distinct mutations were identified, resulting in H147R (three patients), E143K or A131V substitutions in the CDO1 protein **(Fig. 7D)**. Clinical information was reported for three patients (one with each mutation): overlapping features including significant microcephaly, encephalopathy, EEG abnormalities or seizures and hypertonia or abnormal movement^24^. Intriguingly, our genetic and biochemical data highlights both E143 and H147 as critical residues responsible for the interaction between CDO1 and LRRC58 (**Fig. 3 and Fig. S3)**; furthermore, our saturation mutagenesis stability profiling of CDO1 suggests that both the E143K and H147R mutants are refractory to LRRC58-mediated degradation **(Fig. 3A)**. We validated these screen results through individual experiments: both the E143K and H147R CDO1 mutants remained maximally stable even in the absence of extracellular cystine **(Fig. 7E)** and were unable to interact with LRRC58 **(Fig. 7F)**. In contrast, A131V was not identified as a substitution affecting CDO1 stability in our saturation mutagenesis screen **(Fig. 3A)** and this variant behaved as the wild-type protein in our individual validation experiments **(Fig. 7E-F)**. Thus, in light of the lack of functional impairment coupled with the conservative nature of the amino acid substitution, we speculate that A131V may represent a benign variant that is not responsible for the neurodevelopmental abnormalities observed in one patient.

Next we examined the functional impact of the disease mutations. Consistent with the data above, HEK-293T cells expressing either the E143K or H147R CDO1 mutants were unable to survive overnight cystine deprivation, whereas no toxicity was apparent upon expression of either the wild-type protein or the A131V mutant which remain subject to regulation by LRRC58 **(Fig. 7G)**. Cell death occurred by ferroptosis, as it was accompanied by a dramatic increase in lipid peroxidation **(Fig. 7H and Fig. S6C)** and was ameliorated through the addition of Ferrostatin-1 **(Fig. 7G)**. Finally, we considered the effect of CDO1 disease mutants on the differentiation of neural progenitor cells. We established primary neurosphere cultures from fetal brain tissues derived from two independent donors (9 and 12 post conception weeks), wherein neural progenitors are maintained in a state of self-renewal. The neural stem cells were transduced with lentiviral vectors to express CDO1 variants, and then we examined their ability to differentiate into TUJ1^+^ neurons upon serum stimulation over ten days in two-dimensional culture **(Fig. 7I)**. Expression of the E143K and H147R CDO1 disease mutants resulted in a significant reduction in the number of TUJ1^+^ cells generated compared to the wild-type control **(Fig. 7J-K)**, supporting the notion that the toxic effect of these mutations could impair neuronal development. Altogether, these data suggest that loss of LRRC58-mediated regulation underpins the neurodevelopmental defects resulting from CDO1 mutations.

## DISCUSSION

The mechanisms through which cells sense and respond to fluctuations in amino acid abundance remain incompletely defined. Here we demonstrate a role for conditional protein degradation via the ubiquitin-proteasome system in cysteine homeostasis: the Cul2^LRRC58^ E3 ligase complex targets CDO1 for degradation when cysteine is scarce to inhibit cysteine catabolism. Cysteine-dependent regulation occurs at the level of LRRC58 stability, with LRRC58 subject to auto-ubiquitination and degradation only when cysteine is abundant. Interfering with the LRRC58-dependent degradation of CDO1 promotes ferroptosis when cysteine is scarce, and, underscoring the physiological importance of this relationship, CDO1 mutations responsible for severe neurodevelopmental defects encode dominant-active proteins impervious to LRRC58-mediated degradation. Thus, the CDO1-LRRC58 axis is an essential regulator of cysteine homeostasis implicated in neural development.

Our comparative stability profiling screen identified conditional regulation of CDO1 on the basis of differential stability between Jurkat and HEK-293T cells. The cysteine-dependent regulation of CDO1 stability via LRRC58 occurred similarly in both cell types, however, and so we speculate that under typical culture conditions the intracellular cysteine concentration in Jurkat may simply be lower than that in HEK-293T. Interestingly, Cul2^LRRC58^ is active in both cell types despite neither expressing CDO1 endogenously; indeed, RNA-seq data suggests that LRRC58 is uniformly expressed across tissues **(Fig. S7A)**, whereas CDO1 expression is restricted primarily to the liver and a handful of other cell types^35,44^. This discrepancy raises the possibility that the cell may exploit the Cul2^LRRC58^ complex to confer cysteine-sensitive regulation on other pathways, and thus the identification of additional LRRC58 substrates may illuminate other important regulators of cysteine homeostasis.

The activity of LRRC58 is inhibited when cysteine is replete, permitting CDO1 to initiate the conversion of excess cysteine into taurine. Our data demonstrates that conditionality is imparted at the level of LRRC58 abundance, as, when the cysteine-induced auto-degradation of LRRC58 is inhibited (through the K111R mutation, for example), LRRC58 is capable of efficient CDO1 degradation in the presence of cysteine. Self-regulation has been reported previously for several other E3 ubiquitin ligases^45^ including MARCHF6, whose auto-degradation is regulated by cholesterol^46^. Determining the molecular mechanism through which the behavior of LRRC58 is toggled in response to cysteine abundance will be a priority for further study. One possibility is that a hitherto unidentified upstream factor signals cysteine abundance to LRRC58, for example through a post-translational modification; alternatively, LRRC58 itself may bind cysteine and thus act as a direct “sensor” of cysteine concentration. Our saturation mutagenesis profiling data highlights a critical role for residues in the C-terminal domain of LRRC58, and thus one possibility is that cysteine binding in this region results in a conformational change that promotes auto-ubiquitination of LRRC58 via the Cul2 complex. Ultimately, structural insight into the architecture of the Cul2^LRRC58^ complex in the presence and absence of cysteine may be essential to answer this question.

The physiological importance of the CDO1-LRRC58 relationship is underscored by the recent identification of CDO1 mutations as the cause of neurodevelopmental abnormalities including severe microcephaly and encephalopathy^24^. Here we reveal the biochemical basis of this effect, demonstrating that both the E143K and H147R mutations disrupt the interaction interface with LRRC58 and thus permit unrestrained CDO1 activity. The impaired neuron differentiation that we observed *in vitro* coupled with the severe microcephaly observed in patients suggests an important role for the CDO1-LRRC58 axis in safeguarding neuron differentiation; whether this regulation is important for neural progenitor proliferation or the survival of the newly differentiated neurons remains to be determined. One potential implication is that cysteine levels are limiting in the developing brain, an organ known to be particularly sensitive to redox stress^47^, and hence that tight regulation of CDO1 activity is essential to prevent cysteine starvation. Our findings suggest that it may be possible to address this question by developing CDO1 or LRRC58 fusion constructs analogous to those described here to act as biosensors reporting on cysteine abundance in live tissues.

During the final stages of the preparation of this manuscript, our findings were corroborated by an independent study reporting that LRRC58 mediates CDO1 degradation to regulate cysteine catabolism^48^. Examining the role of LRRC58 *in vivo*, the authors show that adeno-associated virus-mediated knockdown of LRRC58 in liver drives increased flux from cysteine to taurine and remodels fatty acid metabolism, suggesting that inhibition of LRRC58 activity may be an effective approach to lower hepatic cholesterol^48^. Cancer therapy is another setting where reducing intracellular cysteine levels may be beneficial, as, although cysteine is considered a non-essential amino acid, a source of exogenous cystine can become essential to drive oncogenic proliferation and mitigate against elevated oxidative stress in tumours^49^. In particular, TCGA data indicates that both high CDO1 expression and low LRRC58 expression are correlated with patient survival in hepatocellular carcinoma **(Fig. S7B)**. Thus, LRRC58 could represent a pharmacological target for the therapeutic manipulation of cysteine levels.

## ACKNOWLEDGEMENTS

We are grateful to Gabriela Grondys-Kotarba and Reiner Schulte at the Cambridge Institute for Medical Research Flow Cytometry Core Facility and to the NIHR Cambridge BRC Cell Phenotyping Hub. We thank Roger Baker, Xiaoling He and the Cambridge Brain Repair Centre for sharing human fetal tissue. This work was supported by an Academy of Medical Sciences Springboard Grant (SBF007\100019), an Isaac Newton Trust/Wellcome ISSF/University of Cambridge Joint Research Grant and an ERC Starting Grant (ERC-2024-STG 101160971) to R.T.T. R.T.T. is a Pemberton-Trinity Fellow. L.H.E.W. was supported by a Postdoctoral Fellowship from the Swedish Research Council (2024-00197) and the IF:Stiftelse, a Swedish foundation for pharmaceutical research, C.C. was supported by a PhD studentship from the UK Medical Research Council (MRC) under the Industrial Cooperative Awards in Science & Technology (iCASE award with Tocris Bio-Techne) doctoral training programme (MR/R015791/1) and is currently a postdoctoral researcher funded by the Michael J. Fox Foundation, and J.B.C. is a recipient of a University of Cambridge School of Biological Sciences DTP PhD Studentship and Peter and Emma Thomsen’s Scholarship (1051). I.A.T. is supported by Wellcome (092096) and Cancer Research UK (C6946/A14492); M.P.W. is supported by a Wellcome Discovery Award (309425/Z/24/Z); N.S.B. is supported by seed funding from the Cambridge Stem Cell Institute, the Royal Society (RGS\R1\231143) and a Wellcome Career Development Award (227294/Z/23/Z), and research led by A.C. on Cullin-RING E3 ligases receives funding from the Innovative Medicines Initiative 2 (IMI2) Joint Undertaking under grant agreement 875510 (EUbOPEN project), which receives support from the European Union’s Horizon 2020 research and innovation program, EFPIA companies and Associated Partners: KTH, OICR, Diamond and McGill.

## AUTHOR CONTRIBUTIONS

D.E.R. and R.T.T. conceived the study. D.E.R., L.H.E.W., C.C., J.B.C., T.A.vW., D.W.G., Z.B., M.A.K., K.H. and M.D. performed the experiments and analyzed the data, supervised by M.P.W, N.S.B., A.C. and R.T.T.. I.A.T. provided essential reagents. R.T.T. wrote the manuscript.

## DATA AVAILABILITY

Additional data and/or reagents that support the findings of this study are available from the corresponding author upon reasonable request.

## DECLARATION OF INTERESTS

A.C. receives or has received sponsored research funding from Almirall, Amgen, Amphista Therapeutics, Boehringer Ingelheim, Eisai, GlaxoSmithKline, Merck KGaA, Nurix Therapeutics, Ono Pharmaceutical and Tocris (a Bio-Techne brand). A.C. is a scientific founder and shareholder of Amphista Therapeutics, a company that is developing targeted protein degradation therapeutic platforms. A.C. is on the Scientific Advisory Board of ProtOS Bio and TRIMTECH Therapeutics.

## MATERIALS & METHODS

### Cell culture

HEK-293T cells were grown in Dulbecco’s Modified Eagle’s Medium (DMEM) (Merck, #D6429) and Jurkat cells were grown in Roswell Park Memorial Institute Medium (RPMI) (Merck, #R8758), both supplemented with 10% fetal bovine serum (Gibco, #A5256701) plus 1% penicillin and streptomycin (Gibco, #15140122). Cells were incubated at 37°C plus 5% CO_2_. Cystine-free DMEM was prepared by adding 200 µM L-glutamine (Sigma-Alrich, #G8540) and 200 µM L-methionine (ThermoFisher Scientific, #63-68-3) to DMEM containing no glutamine, methionine or cystine (Gibco, #21013024).

### Primary human neurosphere culture

Human fetal brain tissue was obtained from the Cambridge University Hospitals NHS Foundation Trust under permission from the NHS Research Ethical Committee (96/085) via the Cambridge Centre for Brain Repair. Two independent primary cultures were established for this study: BRC 2742 (9 post conception weeks) and BRC 2748 (12 post conception weeks). Primary neurosphere cultures were established and maintained as previously described^50^. Briefly, brain tissue was dissociated into single cells using Accutase (ThermoFisher, #A1110501) at 37°C. The cells were filtered (Merck, #CLS352340), pelleted by centrifugation (500 x *g*, 5 min), resuspended in neural stem cell (NSC) media comprised of Neurobasal-A (ThermoFisher Scientific, #10888022) supplemented with 1X B27 (ThermoFisher Scientific, #A3353501), 1X N2 (ThermoFisher Scientific, #11520536), 2 mM L-glutamine (ThermoFisher Scientific, #A2916801), 1X non-essential amino acids (ThermoFisher Scientific, #11140035), 100 U/mL penicillin-streptomycin (ThermoFisher Scientific, #15140122), 20 ng/ml EGF (QKINE, #Qk011-0100) and FGF2 (QKINE, #Qk027-0100), and plated in ultra-low attachment plates (Appleton Woods, #CC227). Growth factors were replenished every other day, and the media were changed weekly. Neurobasal media contains 260 µM L-cysteine.

### Chemicals

The following inhibitors were used: Bortezomib (Cambridge Bioscience, #CAY10008822; used at a final concentration of 1 µM), TAK-243 (Cambridge Bioscience, #HY-100487; 1 µM), MLN4924 (Cambridge Bioscience, #CAY15217; 1 µM), Bafilomycin A1 (ThermoFisher Scientific, #J61835.MCR; 1 µM), Erastin (Cambridge Bioscience, #CAY17754; 1 µM), Sulfasalazine (ThermoFisher Scientific, #461241000; 500 µM), Torin-1 (Stratech Scientific, #A8312-APE; 250 nM), buthionine sulfoximine (BSO) (Merck, #19176; 2 µM) and Ferrostatin-1 (Merck, #SML0583; 1 µM). Hydrogen peroxide was obtained from Fisher Scientific (#10386643; 100 µM), glutathione ethyl ester was obtained from Cambridge Bioscience (#CAY14953; 500 µM) and DTT was obtained from ThermoFisher Scientific (#R0861; 50 µM).

### Antibodies

Primary antibodies used were: rabbit anti-CDO1 (Proteintech, #12589-1-AP), rabbit anti-GFP (Abcam, #ab290), mouse anti-FLAG M2 (Merck, #F3165), rabbit anti-HA C29F4 (Cell Signaling #3724), mouse anti-Ubiquitin (Life Sensors, #VU101), rabbit anti-Cul2 (Bethyl Laboratories, #A302-476A), rabbit anti-Cul5 (Bethyl Laboratories, #A302-173A), mouse anti-Vinculin (Sigma, #V9131), mouse anti-β-actin (Sigma, #A2228) and rabbit anti-TUJ1 (Abcam, #ab18207). Secondary antibodies used were: Alexa Fluor Plus 555-conugated donkey anti-rabbit IgG (ThermoFisher Scientific, #A32794), HRP-conjugated donkey anti-mouse IgG (Jackson ImmunoResearch #715-035-150) and HRP-conjugated donkey anti-rabbit IgG (Jackson ImmunoResearch #711-035-152).

### Plasmids

Stability profiling experiments were performed using the pHAGE-GPS lentiviral vector, with either CDO1 or LRRC58 cloned between the BstBI and XhoI sites to generate a C-terminal fusion to GFP. Lentiviral expression of LRRC58 was achieved using either pHRSIN-based lentiviral vectors (pHRSIN-P_SFFV_-LRRC58-WPRE-P_PGK_-Hygro^R^) or pKLV2-based lentiviral vectors (pKLV2-P_SFFV_-LRRC58-P_PGK_-Puro^R^-P2A-BFP-WPRE). CDO1 expression was either achieved using the pHAGE-GPS vector described above, or, for experiments involving C11-BODIPY staining, a pKLV2 vector (pKLV2-P_SFFV_-CDO1-P_PGK_-Puro^R^-P2A-BFP-WPRE). All molecular cloning was performed using the Gibson assembly method (NEBuilder HiFi DNA Assembly Master Mix, NEB #E2621S).

### CRISPR/Cas9-mediated gene disruption

CRISPR sgRNA were synthesized as top and bottom strand oligonucleotides (IDT), phosphorylated using T4 PNK (NEB #M0201), and annealed by heating at 95°C for 5 min followed by cooling to room temperature at 0.1°C per min. T4 ligase (NEB #M0202) was then used to ligate the fragments into lentiCRISPRv2 (Addgene #52961, kindly deposited by Feng Zhang^51^) cut with BsmBI (NEB #R0739S). Targeting sequences used were:

sg1-Control (targets an intron in *FOXP1*): gTGGGAACAGGATGAGGAAGG

sg2-Control (targets an intron in *ATP1A1*): GATGGGCAAGAAGGAAGCAG

sg1-*LRRC58*: gCCGTGCGGTGGCCAGGATGG

sg2-*LRRC58*: gTGCGGTGGCCAGGATGGAGG

sg3-*LRRC58*: gCCCGTGGCAGCGACACCAGA

sg4-*LRRC58*: GCTGCTGCCTCACAACCGTC

sg*CUL2*: gTTTGACGACAATAAAAGCCG

sg1-*CUL5*: gCCTAGCTTATGCTTGATACA

sg2-*CUL5*: GGAATGCTGTGTAAATGCCC

### Lentivirus production

HEK-293T cells at ∼80% confluency were transfected with the lentiviral transfer vector of interest in addition to a viral packaging mix containing four plasmids encoding Gag-Pol, Rev, Tat and VSV-G using PolyJet In Vitro DNA Transfection Reagent (SignaGen Laboratories, #SL100688). The manufacturer’s protocol was followed, with the exception that the media was not replaced prior to transfection. The next morning the media was replaced, and then the lentiviral supernatant was collected 48 hours post-transfection. Cell debris was removed either by centrifugation or using a 0.45 μm filter, and the resulting lentiviral stocks were either used immediately or stored in single-use aliquots at - 80°C.

### Flow cytometry and FACS

Flow cytometry was carried out on an LSR-II instrument (BD Biosciences) using FACS Diva software. A minimum of 10,000 live cells were collected per condition. The resulting data was analyzed using FlowJo (version 10.4). For FACS, cells were washed twice in PBS, resuspended in sort solution (PBS plus 2% FBS) and passed through a 50 µm filter (Sysmex, #04-004-2327). FACS was carried out using an Influx Cell Sorter (BD Biosciences), with sorted cells recovered in collection media comprised of a 1:1 mixture of complete media and FBS.

### Protein structure prediction

Protein structure predictions were generated using AlphaFold 3 run via the AlphaFold Server (https://alphafoldserver.com/) and subsequently analyzed using UCSF ChimeraX (version 1.9).

### GPS stability profiling

Two modifications were made to the existing barcoded GPS-uORFeome expression library^22^ to facilitate expression in Jurkat cells. First, the human cytomegalovirus promoter (P_CMV_) was replaced with the Rous Sarcoma Virus promoter (P_RSV_), which was achieved by excising the P_CMV_-DsRed-IRES-GFP cassette through digestion with the PI-SceI (NEB, #R0696S) and I-PpoI (Promega, #R7031) restriction enzymes and replacing it with an P_RSV_-DsRed-IRES-GFP cassette amplified by PCR (Q5 Hot Start High-Fidelity DNA Polymerase, NEB #M0493S) using the Gibson assembly method (NEBuilder HiFi DNA Assembly Master Mix, NEB #E2621S). Reaction products were concentrated using SPRIselect beads (Beckman Coulter, #B23317) and electroporated into ElectroMAX DH10B *E. coli* (ThermoFisher Scientific, #18290015) as per the manufacturer’s protocol. The bacteria were recovered in S.O.C. Medium (ThermoFisher Scientific, #15544034), and, following incubation for 1 h at 37°C with shaking, were grown overnight on LB agar plates supplemented with ampicillin (100 μg/ml) at 30°C. Plasmid DNA was extracted from the bacterial lawns using the GenElute HP Plasmid Midiprep Kit (Merck, #NA0200), ensuring that sufficient colonies were recovered to maintain a minimum of 100-fold representation of the library. Second, an analogous cloning process was carried out to remove the P_PGK_-Puro^R^ cassette, which was achieved through digestion with I-SceI (NEB, #R0694S) and PacI (NEB, #R0547S) restriction enzymes.

The resulting P_RSV_-GPS-ORFeome library was packaged into lentiviral particles and introduced into Jurkat cells at a multiplicity of infection of ∼0.3 (∼30% DsRed+ cells), such that the overwhelming majority of cells expressed no more than one single GFP-fusion protein. Four days post-transduction, the population was partitioned into six stability bins based on the stability of the GFP-fusion protein (GFP/DsRed ratio), such that each bin received approximately the same number of cells. Genomic DNA was then extracted from each of the sorted populations using the Gentra Puregene Cell Kit (Qiagen, #158767). To quantify the distribution of each ORF across the six stability bins, the molecular barcodes located at the 3’ end of each ORF were amplified through 24 cycles of PCR (Q5 Hot Start High-Fidelity DNA Polymerase, NEB #M0493S) using primers annealing to invariant flanking sites; a pool of 8 “staggered” forward primers were used to introduce nucleotide diversity for the Illumina platform. Following purification using a spin column (Qiagen QIAquick PCR Purification Kit, **#**28104), 200 ng was used for a second PCR reaction (7 cycles) to add the Illumina P5 and P7 adaptors and indexes for multiplexing. Samples were again purified using a spin column, quantified by Nanodrop spectrophotometry, pooled evenly, and then excised from a 2% agarose gel (Qiagen QIAEX II Gel Extraction Kit, #20021). Sequencing was performed on an Illumina NovaSeq 6000 instrument.

The resulting raw sequence reads were trimmed of constant flanking sequences using Cutadapt (version 4.1) and aligned to a reference index detailing all of the ORF-barcode combinations using Bowtie 2 (version 2.5.4). To reflect the distribution of sequence reads across the six stability bins, a protein stability index (PSI) metric was calculated given by the sum of multiplying the proportion of reads in each bin by the bin number, resulting in a stability score between 1 (maximally unstable) and 6 (maximally stable). To facilitate comparison between the PSI scores generated from a previous stability profiling screen carried out in HEK-293T cells^25^, scores were rescaled between 0 (maximally unstable) and 1 (maximally stable) via min-max normalization and used to calculate a ΔPSI metric **(Table S1)**.

### CRISPR/Cas9 genetic screens

Using the lentiCRISPRv2 dual sgRNA/Cas9 expression vector^22^, we previously constructed a ubiquitin-focused sgRNA library comprising 6 guides per gene targeting ∼1500 genes with a known or postulated functional role within the ubiquitin-proteasome system^22^. We packaged this library into lentiviral particles and introduced it into HEK-293T cells expressing either GPS-CDO1 or GPS-LRRC58 at single copy. Untransduced cells were eliminated through two rounds of selection with puromycin (1.5 µg/ml), each for two days. Seven days post-transduction, cells exhibiting stabilization of the GFP fusion constructs were isolated by FACS, targeting the top ∼3% of cells displaying the highest GFP/DsRed ratio; in the case of the GPS-CDO1 screen, cystine-free media was applied to the cells the day before the sort. Genomic DNA was extracted from both the sorted cells (Zymo Quick-DNA Microprep Kit, #D3021) and the unsorted library (Qiagen Gentra Puregene Cell Kit, #158767). The sgRNAs in each population were amplified by two rounds of PCR in a manner analogous to that described above and prepared for Illumina sequencing in the same way. The resulting sequence reads were trimmed of constant flanking sequences using Cutadapt (version 4.1) and aligned to a reference index detailing all of the sgRNAs in the library using Bowtie 2 (version 2.5.4). The resulting raw count table was analyzed using the MAGeCK algorithm^52^ to identify genes targeted by multiple sgRNAs enriched in the sorted population.

### Saturation mutagenesis stability profiling of CDO1

The 200 amino acids comprising CDO1 were considered in four segments of equal size: 1-50, 51-100, 101-150 and 151-200. For each segment, an oligonucleotide pool was designed such that each residue was substituted with all other possible residues; to maximize sequence diversity between library members, codons were assigned via a weighted random selection process based on codon usage frequency across the human proteome. The oligonucleotide pool was synthesized (Twist Bioscience), and each mutagenic segment amplified by PCR using primers binding to common flanking regions. These flanking regions also served as sites of overlap for a subsequent Gibson assembly reaction. The PCR products were purified via extraction from a 2% agarose gel (QIAEX II gel extraction kit, QIAGEN #20021) and then assembled with the GPS lentiviral vector cut BstBI (NEB, #R0519S) and XhoI (NEB, #R0146S) along with the corresponding wild-type fragment(s) of CDO1 required to reconstitute full-length CDO1; for example, mutant segment 1 was assembled with a fragment encoding wild-type segments 2-4, and mutant segment 2 was assembled with two other fragments, one encoding wild-type segment 1 and the other encoding wild-type segments 3-4. The assembled products were electroporated into DH10B *E. coli* and the resulting plasmid DNA purified as described above.

Each library was packaged into lentiviral particles and introduced at single copy into wild-type HEK-293T cells in duplicate. Following puromycin selection to eliminate untransduced cells, the resulting populations were depleted of cystine overnight and then partitioned by FACS into four stability bins based on the stability (GFP/DsRed ratio) of the mutant CDO1 construct. The sorted cells were processed as described above and the abundance of each variant across the four sequencing bins quantified by PCR and Illumina sequencing (150 bp paired-end reads). Bioinformatic analysis of the resulting sequence reads was performed in the same way to that described above, yielding a PSI metric between 1 (maximally unstable) and 4 (maximally stable) for each CDO1 variant. For the heatmaps displayed in Figure 3, the mean of the two duplicate PSI scores is shown with the exception of segment 4 where, owing to poor concordance between the two replicates, the data for replicate 1 was discarded in favor of replicate 2 alone.

### Saturation mutagenesis profiling of LRRC58

Residues 156-371 of LRRC58 were considered as 4 segments for mutagenesis, each comprising 54 amino acids: 156-209, 210-263, 264-317 and 318-371. The saturation mutagenesis library was designed and amplified by PCR in an equivalent manner to that described above. However, in contrast to the multi-fragment Gibson assembly cloning approach used for the CDO1 saturation mutagenesis library, here we first created a series of intermediate vectors to simplify the cloning process. Starting with a modified version of the pKLV2 lentiviral vector (Addgene #67974, kindly deposited by Kosuke Yusa^53^) encoding LRRC58 downstream of the SFFV promoter, we used Gibson assembly to create four “stuffer” vectors in which the sequence encoding the region to be mutagenized was replaced with a short “stuffer” sequence which could be excised via flanking BbsI restriction sites. Thereafter, these vectors were cut with BbsI (NEB, #R3539S), purified using SPRI beads, and mixed with the PCR product encoding the corresponding mutant segment amplified from the oligonucleotide pool in a Gibson assembly reaction and subsequently processed as described above. These libraries were used directly for the LRRC58 activity profiling experiment; for the LRRC8 stability profiling experiment, the four mutant libraries were first subcloned into the GPS expression vector.

Each of the eight libraries was packaged into lentiviral particles and introduced at single copy into either wild-type HEK-293T cells (stability profiling) or a LRRC58 KO HEK-293T clone expressing GPS-CDO1 (activity profiling) in duplicate. Following puromycin selection to eliminate untransduced cells, the resulting populations were partitioned by FACS into three stability bins based on the stability (GFP/DsRed ratio) of either the GPS-LRRC58 construct (stability profiling) or the GPS-CDO1 construct (activity profiling). The sorted cells were processed as described above and the abundance of each variant across the three sequencing bins quantified by PCR and Illumina sequencing (150 bp paired-end reads). Bioinformatic analysis of the resulting sequence reads was performed in the same way to that described above, yielding a PSI metric between 1 (maximally unstable) and 3 (maximally stable) for each LRRC58 variant.

### Immunoblotting

Cells were harvested, washed once in PBS, and lysed in 1% SDS plus 1:200 Benzonase (Merck, #E1014) for 20 minutes at room temperature. 4X Laemmli Sample Buffer (Bio-Rad, #1610747) was then added and the lysates were heated to 70°C for 10 min. SDS-PAGE was performed using 4-12% Bis-Tris gels (Merck, #MP41G12). Proteins were transferred onto PVDF membrane (Merck, #IPVH00010) activated in methanol using the Trans-Blot SD Semi-Dry Transfer System (Bio-Rad). For the visualization of ubiquitinated LRRC58 using the VU-1 antibody, membranes were first cross-linked with 0.5% glutaraldehyde (ThermoFisher Scientific, #111-30-8); otherwise, membranes were immediately blocked for a minimum of 30 min in 5% Skim Milk Powder (Merck, #70166) dissolved in PBS supplemented with 0.2% Tween-20 (PBS-T) (Sigma-Aldrich, #P1379). Membranes were subsequently incubated with primary antibodies overnight at 4°C, washed at least three times in PBS-T, and then incubated with HRP-conjugated secondary antibodies for 40 min at room temperature. Following a further five washes in PBS-T, reactive bands were visualized using SuperSignal West Detection Reagents ECL (Pierce, #32106), Pico (Pierce, #34580) or Dura (Pierce, #34075). Images were collected on a ChemiDoc Imaging System (Bio-Rad) and further processed using GNU Image Manipulation Platform (GIMP) version 2.10.34.

### Immunoprecipitation

Cells were harvested, washed once in ice-cold PBS, and then lysed in 1% IGEPAL (Sigma-Aldrich) dissolved in TBS supplemented with an EDTA-free protease inhibitor cocktail tablet (Roche, #11697498001) on ice for 20 minutes. Insoluble material was pelleted by centrifugation (21,000 x *g*, 10 min, 4°C); 5 µl ChromoTek GFP-Trap Magnetic Agarose beads (Proteintech, #gtma) were then added to the soluble fraction and the samples incubated for 2 h at 4°C with continuous rotation. Following three 5-minute washes with lysis buffer, samples were eluted in 1X Laemmli Sample Buffer (Bio-Rad, #1610747) and analyzed by immunoblot as described above.

### Validation of knockout clones generated through CRISPR/Cas9-mediated gene disruption

Genomic DNA was extracted using the Gentra Puregene Cell Kit (Qiagen, #158767). The sequence encoding exon 1 of LRRC58 was amplified by PCR (Q5 Hot Start High-Fidelity DNA Polymerase, NEB #M0493S); following spin column purification, a second round of PCR was performed to attach Illumina P5 and P7 adaptors and indexes for multiplexing. Samples were again purified using a spin column, quantified by Nanodrop spectrophotometry, pooled evenly and sequenced on an Illumina NovaSeq 6000 instrument (150 bp paired-end reads). The resulting sequence reads were analyzed for CRISRP/Cas9-induced indels using CRISPResso2 ^54^.

### TMT whole cell proteomics

Approximately 10 million HEK-293T cells were cultured in triplicate in either complete media or media lacking cystine and methionine for 12 h. The cells were then harvested in ice-cold PBS and washed once with ice-cold PBS. Cells were lysed through the addition of a lysis buffer composed of 6 M guanidine HCl (ThermoFisher Scientific, #24115) and 50 mM HEPES (Merck, #H0887) for 10 min at room temperature. The lysates were then sonicated, before cell debris was removed by centrifugation (21000 x *g*, 10 min, 4°C). Dithiothreitol (DTT) was then added at a final concentration of 5 mM and the samples incubated at room temperature for 20 min. Alkylation of cysteine residues was then achieved through the addition of 15 mM iodoacetamide followed by incubation for 20 min at room temperature in the dark. Excess iodoacetamide was quenched with DTT for 15 min. The samples were then diluted by adding 200 mM HEPES pH 8.5 to reach a guanidine concentration of 1.5 M, allowing for digestion with LysC protease (1:100 protease-to-protein ratio) at room temperature for 3 h. The samples were then diluted again with 200 mM HEPES pH 8.5 to reach a guanidine concentration of 0.5 M, allowing for digestion with trypsin (1:100 protease-to-protein ratio) overnight at 37°C. The reaction was quenched through the addition of 5% formic acid and then undigested protein was removed by centrifuged (21000 x *g*, 10 min, 4°C). The resulting peptides were then subjected to C18 solid-phase extraction (Sep-Pak, Waters) and vacuum-centrifuged to near-dryness. Subsequent TMT labeling, MS data acquisition and data analysis were performed exactly as described previously^55^. The TMT labelling strategy was as follows: Complete_rep1, 126; Complete_rep2, 128N; Complete_rep3, 129C; Depleted_rep1, 127N; Depleted_rep2, 128C; Depleted_rep3, 130N.

### Crystal violet staining

Cells were harvested into 5 ml round bottom tubes, washed twice with 3 ml PBS, and then fixed through the addition of 250 µl 100% ethanol for 20 min at 4°C. The samples were washed again with 3 ml PBS and then cellular material was stained by adding 100 µl 1X Crystal Violet solution (Pro-Lab, #PL.7000) for 30 minutes at room temperature. The samples were then washed twice with 3 ml PBS. Finally, bound dye was solubilized by adding 150 µl 100% ethanol and the resulting solution was transferred into a 96-flat bottom plate. The absorbance at 595 nm was measured with the lid removed using a CLARIOstar Plus instrument (BMG Labtech).

### C11-BODIPY staining

.Lipid peroxidation was assessed by adding C11-BODIPY dye (ThermoFisher Scientific, #D3861) into the cell culture media at a final concentration of 2 µM, followed by incubation at 37°C for 30 min. The cells were then harvested and analyzed by flow cytometry as described above.

### *In vitro* differentiation of primary human neural stem cells

Lentiviral supernatants were passed through a 0.45 µm filter and then concentrated 20-fold using Lenti-X Concentrator (Takara, #631231). Primary neurosphere cultures were mechanically dissociated by pipetting and incubated with lentivirus overnight in NSC media supplemented with growth factors and 4 µg/ml protamine sulphate (Merck, #1101230005). The media was changed the next morning, and the cultures were maintained as described above. We observed transduction efficiencies of 30 ± 5% (n = 3, BRC 2742) and 35 ± 8% (n = 3, BRC 2748) as assayed by flow cytometry, with no significant differences between the CDO1 variants examined. *In vitro* differentiation was performed as previously described^50^. Briefly, chamber slides (Thistle Scientific, #IB-80806) were coated with poly-D-lysine (ThermoFisher Scientific, #A389040) and laminin-521 (ThermoFisher Scientific, #A29249) and stored at 4°C. On the day of differentiation, the plates were incubated for >1 h at 37°C. Neurospheres were dissociated into single cells using Accutase, washed and resuspended in NSC media supplemented with 20 ng/mL EGF and FGF2. Cells were plated at a density of 100,000 cells/cm^2^ on the coated plates; the next day, the media was switched to differentiation medium comprised of NSC media plus 2% fetal bovine serum (ThermoFisher Scientific, #10438026). Subsequently the media was changed every other day for 10 days.

### Immunofluorescence microscopy

Cells were fixed at room temperature using 4% paraformaldehyde (ThermoFisher Scientific, #043368.9M) for 15 minutes. Cells were blocked for 1 h at room temperature in blocking buffer, comprising 1X PBS plus 5% bovine serum albumin (Merck, #A9418-50G) and 0.1% Triton X-100 (Sigma Aldrich, #T8787). Primary antibody incubation was performed in blocking buffer overnight at 4°C. Cells were then washed three times with 1X PBS plus 0.1% Triton X-100 for 5 min. Secondary antibody incubation was performed in blocking buffer for 1 hour at room temperature, and samples were protected from light for the rest of the procedure. Cells were washed three times with 1X PBS plus 0.1% Triton X-100 for 5 min and counterstained with DAPI (Sigma-Aldrich, #MBD0015). Images were acquired on an Andor BC43 CF confocal microscope with identical acquisition settings and processed using ImageJ (NIH); quantification was performed manually using the ImageJ/FIJI Cell Counter Plugin (NIH) considering 3 independent images per well which were then averaged.

## SUPPLEMENTARY FIGURE LEGENDS

**Supplementary Figure 1.**
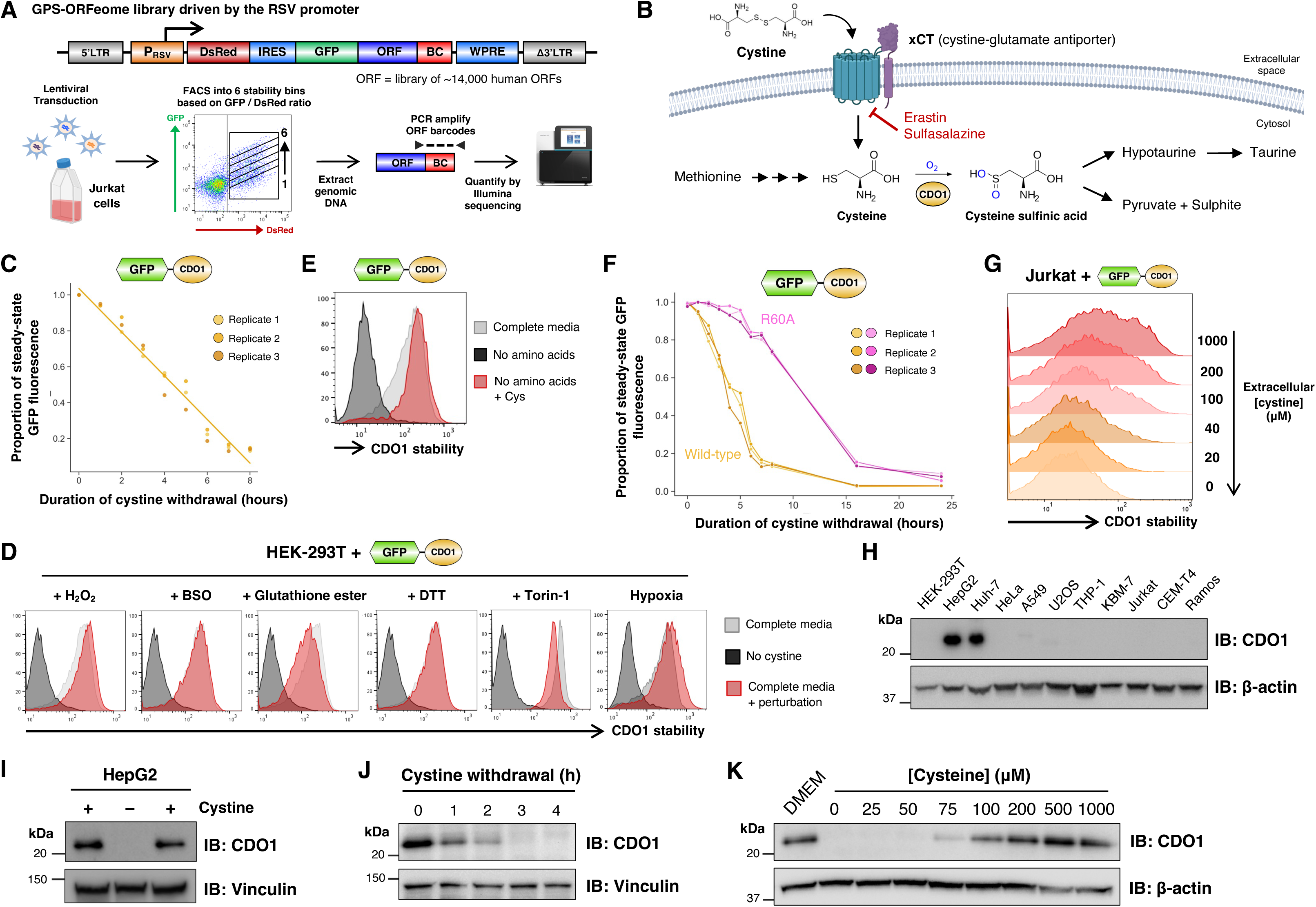
The degradation of CDO1 is cysteine-dependent. **(A)** Schematic representation of the stability profiling screen in Jurkat cells. **(B)** Schematic representation of the role of CDO1 in cysteine metabolism. **(C)** Time course of GPS-CDO1 degradation in HEK-293T cells following cystine removal, as measured by flow cytometry. **(D)** CDO1 degradation is not promoted by mTOR inhibition or redox stress. HEK-293T cells expressing GPS-CDO1 were challenged with the indicated perturbations overnight and then assayed by flow cytometry. **(E)** The degradation of CDO1 induced by total amino acid deprivation can be prevented by re-addition of cystine alone. HEK-293T cells expressing GPS-CDO1 were incubated overnight in complete media, media lacking all amino acids, or media lacking all amino acids except cystine and then analyzed by flow cytometry. **(F)** Lack of CDO1 catalytic activity delays CDO1 degradation following cystine withdrawal. HEK-293T cells expressing either wild-type CDO1 or a catalytically-inactive CDO1 mutant (R60A) in the context of the GPS expression vector were monitored by flow cytometry following cystine withdrawal. **(G)** CDO1 stability is similarly correlated with cysteine abundance in Jurkat. **(H)** CDO1 is expressed by the hepatocellular carcinoma cell lines HepG2 and Huh-7. Expression of endogenous CDO1 protein in the indicated cell lines was assessed by immunoblot. **(I-K)** Endogenous CDO1 abundance is regulated by cysteine levels in HepG2 cells, as assessed by immunoblot. Cystine withdrawal results in loss of CDO1 protein **(I)**; this effect occurs rapidly **(J)** and is correlation with the extracellular cystine concentration **(K)**.

**Supplementary Figure 2.**
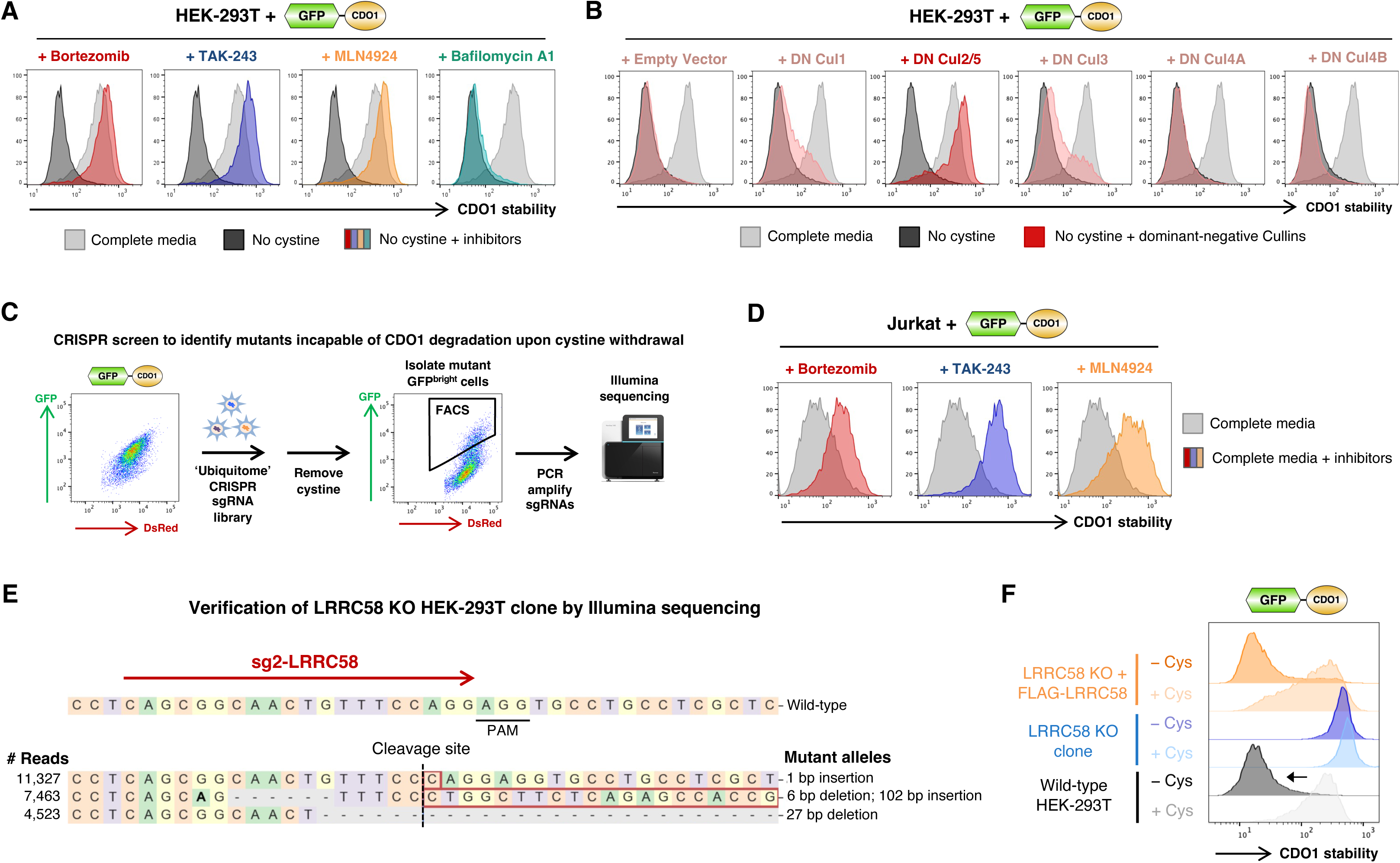
LRRC58 is responsible for the proteasomal degradation of CDO1. **(A)** CDO1 is targeted for ubiquitination and proteasomal degradation by a Cullin-RING E3 ligase complex. HEK-293T cells expressing GPS-CDO1 were treated with the indicated inhibitors in cystine-free media and assessed by flow cytometry 7 hours later. **(B)** CDO1 is degraded by a Cul2 or Cul5 E3 ligase complex. C-terminally truncated dominant-negative Cullin constructs were introduced into HEK-293T cells expressing GPS-CDO1 by lentiviral transduction and CDO1 stability assayed by flow cytometry 48 hours later following cystine withdrawal. **(C)** Schematic representation of the CRISPR screen designed to identify the E3 ligase responsible for CDO1 degradation in the absence of cysteine. **(D)** CDO1 is also targeted for ubiquitination and proteasomal degradation by a Cullin-RING E3 ligase complex in Jurkat. Jurkat cells expressing GPS-CDO1 were treated with the indicated inhibitors and assessed by flow cytometry 7 hours later. **(E)** Validation of a HEK-293T LRRC58 KO clone. LRRC58 exon 1 was amplified by PCR from genomic DNA and CRISPR/Cas9-induced indels characterized by Illumina sequencing. **(F)** Genetic complementation restores CDO1 degradation. The LRRC58 KO clone expressing GPS-CDO1 was transduced with a lentiviral vector encoding FLAG-tagged LRRC58; CDO1 stability was then assessed by flow cytometry in the presence and absence of extracellular cystine.

**Supplementary Figure 3.**
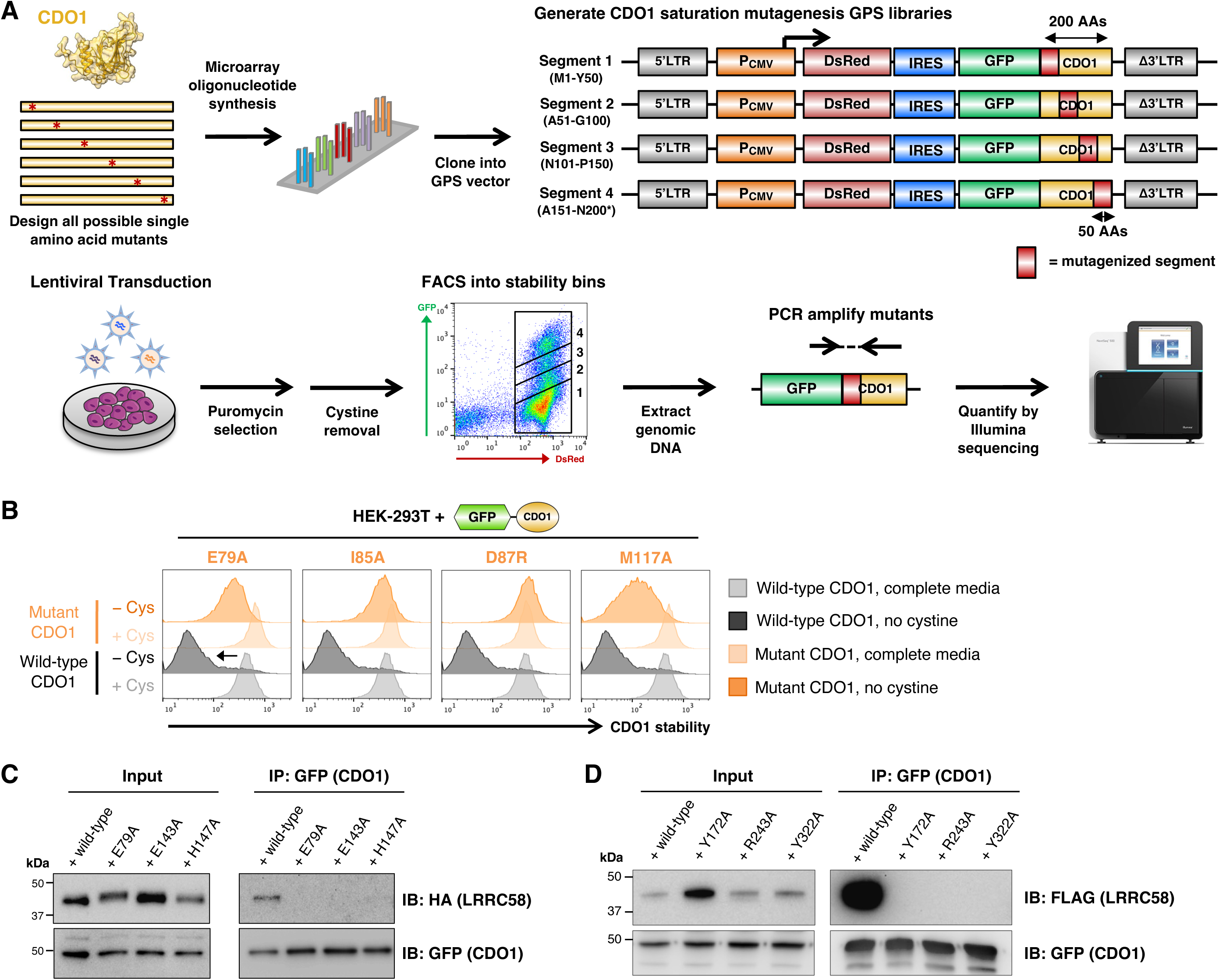
Defining the residues of CDO1 required for cysteine-induced instability. **(A)** Schematic representation of the saturation mutagenesis stability profiling screen designed to identify substitution mutations rendering CDO1 refractory to degradation in the absence of cysteine. Four mutagenic libraries (each covering 50 amino acids) were created in the context of the GPS expression vector, in which each residue of CDO1 was mutated to all other possible amino acids. The resulting libraries were expressed in HEK-293T cells and the stability of each mutant in the absence of extracellular cystine quantified by FACS and Illumina sequencing. **(B)** Individual validation of critical residues required for CDO1 degradation. The indicated CDO1 mutants were expressed in HEK-293T cells in the context of the GPS system, and stability monitored by flow cytometry in the presence and absence of cystine. See also Fig. 3C. **(C)** CDO1 mutants that remain stable upon cystine withdrawal fail to interact with LRRC58. Following overnight treatment with Bortezomib, the indicated CDO1 mutants were immunoprecipitated from HEK-293T cells using an anti-GFP nanobody, and co-immunoprecipitating HA-tagged LRRC58 assessed by immunoblot. **(D)** Mutation of LRRC58 residues lying on the predicted interaction interface abolish the interaction with CDO1. Following overnight treatment with Bortezomib, CDO1 was immunoprecipitated from HEK-293T cells expressing the indicated FLAG-tagged LRRC58 mutants using an anti-GFP nanobody and binding assessed by immunoblot.

**Supplementary Figure 4.**
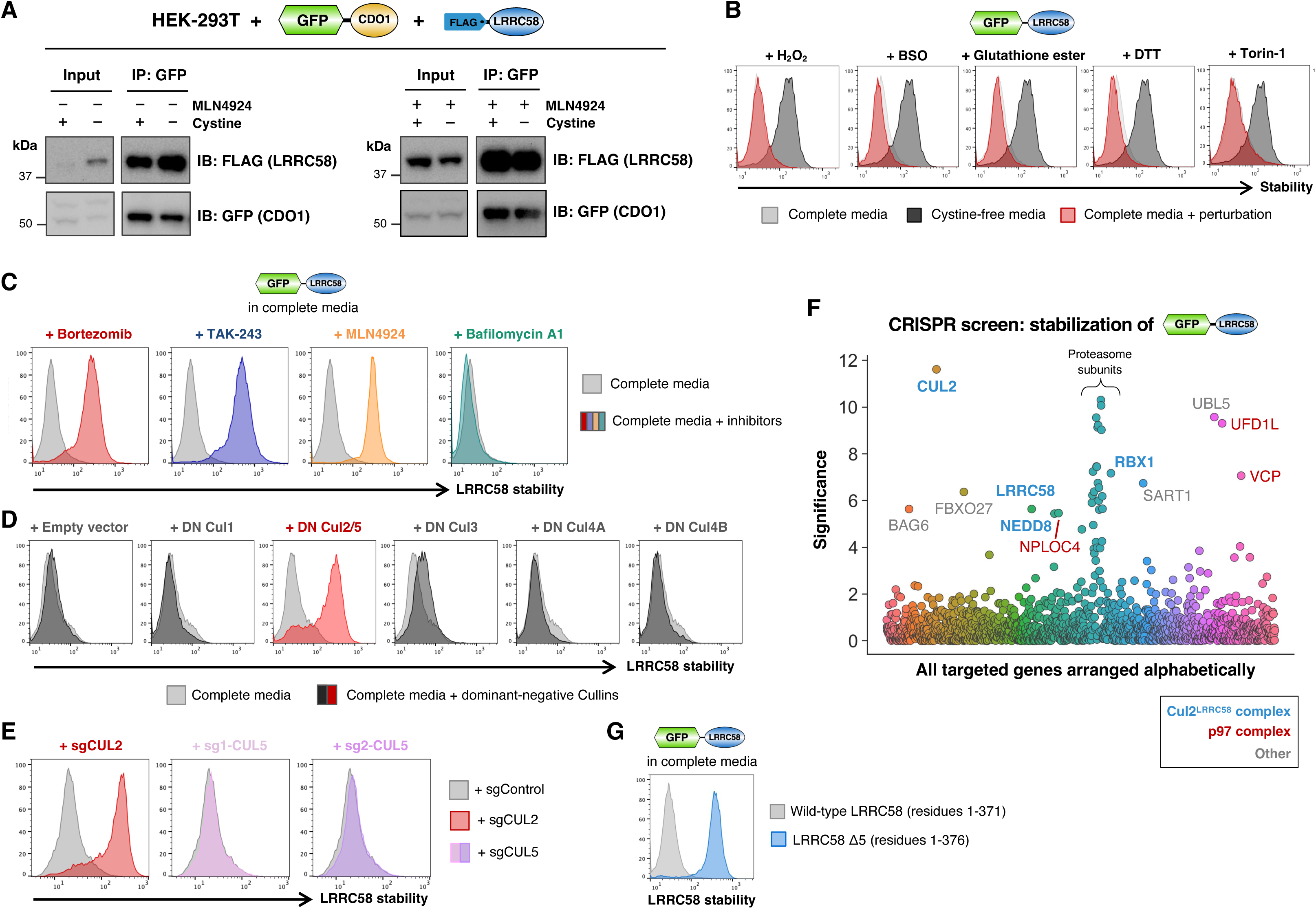
The activity of LRRC58 is restrained by proteasomal degradation. **(A)** CDO1 and LRRC58 interact in both the presence and absence of cysteine. GFP-tagged CDO1 was immunoprecipitated from HEK-293T cells expressing FLAG-tagged LRRC58 in the presence of absence of cystine with and without Cullin-RING ligase inhibition via MLN4924. **(B)** LRRC58 is not stabilized by mTOR inhibition or redox stress. HEK-293T cells expressing GPS-LRRC58 were challenged with the indicated perturbations overnight and then assayed by flow cytometry. **(C-E)** LRRC58 degradation is mediated via a Cul2 E3 ligase complex. **(C)** HEK-293T cells expressing GPS-LRRC58 were challenged with the indicated inhibitors and LRRC58 stabilization assessed by flow cytometry 7 hours later. **(D)** HEK-293T cells expressing GPS-LRRC58 were transduced with lentiviral vectors encoding dominant-negative Cullin constructs and LRRC58 stabilization assessed by flow cytometry two days later. **(E)** CRISPR/Cas9-mediated disruption of *CUL2*, but not *CUL5*, stabilized GPS-LRRC58 in complete media. **(F)** A CRISPR screen to identify the machinery required for the instability of LRRC58 in complete media. HEK-293T cells expressing GPS-LRRC58 were transduced with a CRISPR sgRNA library targeting ubiquitin system components, and mutant GFP^bright^ cells unable to degrade LRRC58 were isolated by FACS. “Significance” on the y-axis represents the negative log of the “pos|score” metric reported by the MAGeCK algorithm^52^. *CUL2* was identified as the leading hit, but no other Cul2 substrate adaptors beyond LRRC58 itself were significantly enriched. **(G)** The extreme C-terminus of LRRC58 is required for cysteine-dependent instability. Wild-type LRRC58 expressed in the context of the GPS system is unstable in complete media, but this effect is abolished upon removal of its last five residues (“Δ5”).

**Supplementary Figure 5.**
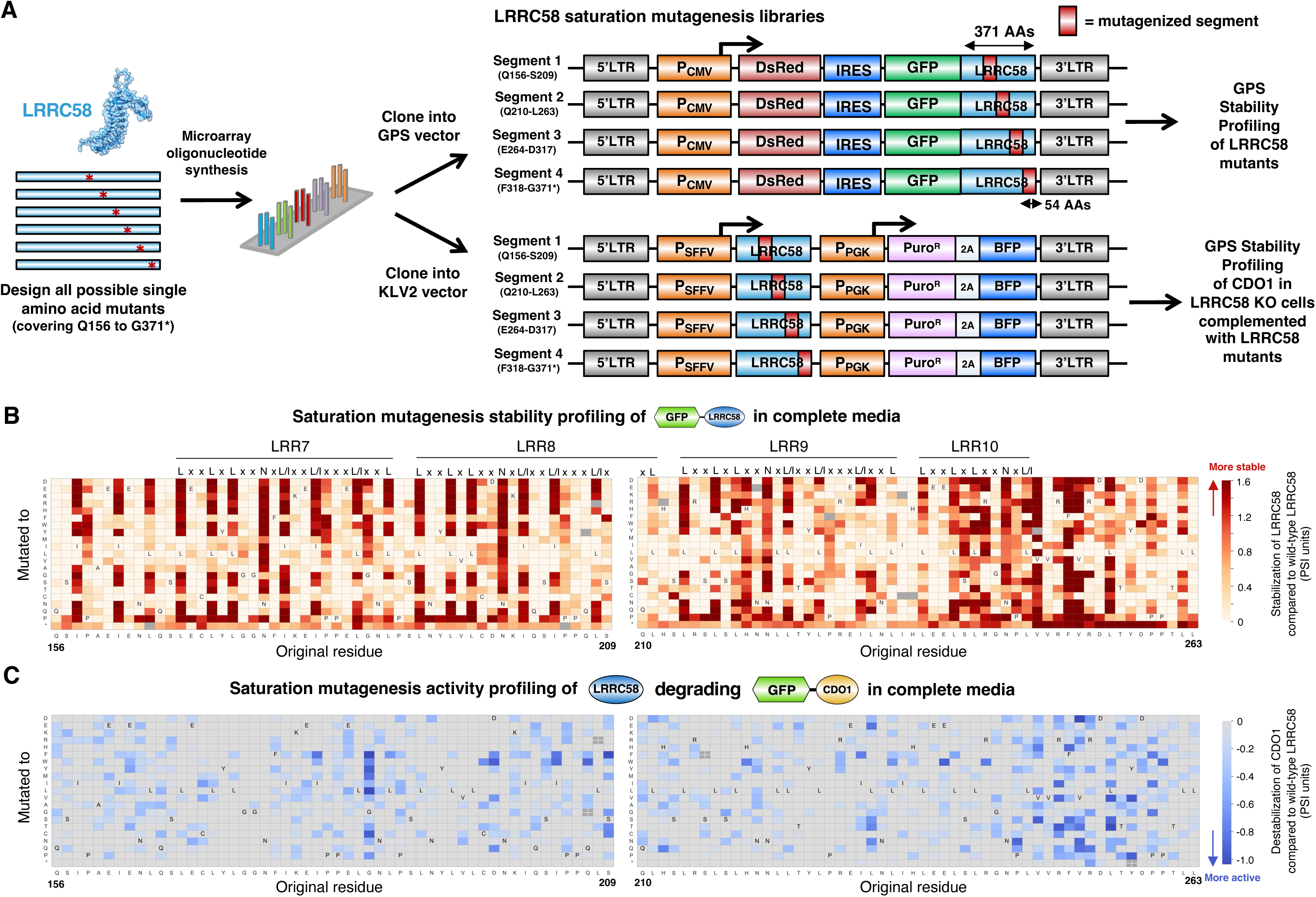
Site-saturation mutagenesis of LRRC58 identifies mutants which hyperactivate CDO1 degradation. **(A)** Schematic representation of the twin LRRC58 saturation mutagenesis screens. Four mutagenic libraries (each covering 54 amino acids) were created in the context of both the GPS expression vector (top) and the KLV2 expression vector (bottom), in which each residue of LRRC58 from Q156 onwards was mutated to all other possible amino acids. The resulting libraries were expressed in either wild-type HEK-293T cells (for stability profiling of LRRC58) or LRRC58 KO cells expressing GPS-CDO1 (for activity profiling of LRRC58). In each case, the performance of each mutant was quantified by FACS and Illumina sequencing. **(B)** Saturation mutagenesis stability profiling of LRRC58 in complete media. Each cell represents the performance of an individual substitution mutant; the darker the red color, the greater the degree of stabilization compared to wild-type LRRC58. Consensus motifs for the leucine-rich repeats are indicated. See also Fig. 5A. **(C)** Saturation mutagenesis activity profiling of LRRC58, in which the ability of LRRC58 mutants to degrade CDO1 was assessed in complete media. Each cell represents the performance of an individual substitution mutant; the darker the blue color, the more hyperactive the mutant compared to wild-type LRRC58. See also Fig. 5B.

**Supplementary Figure 6.**
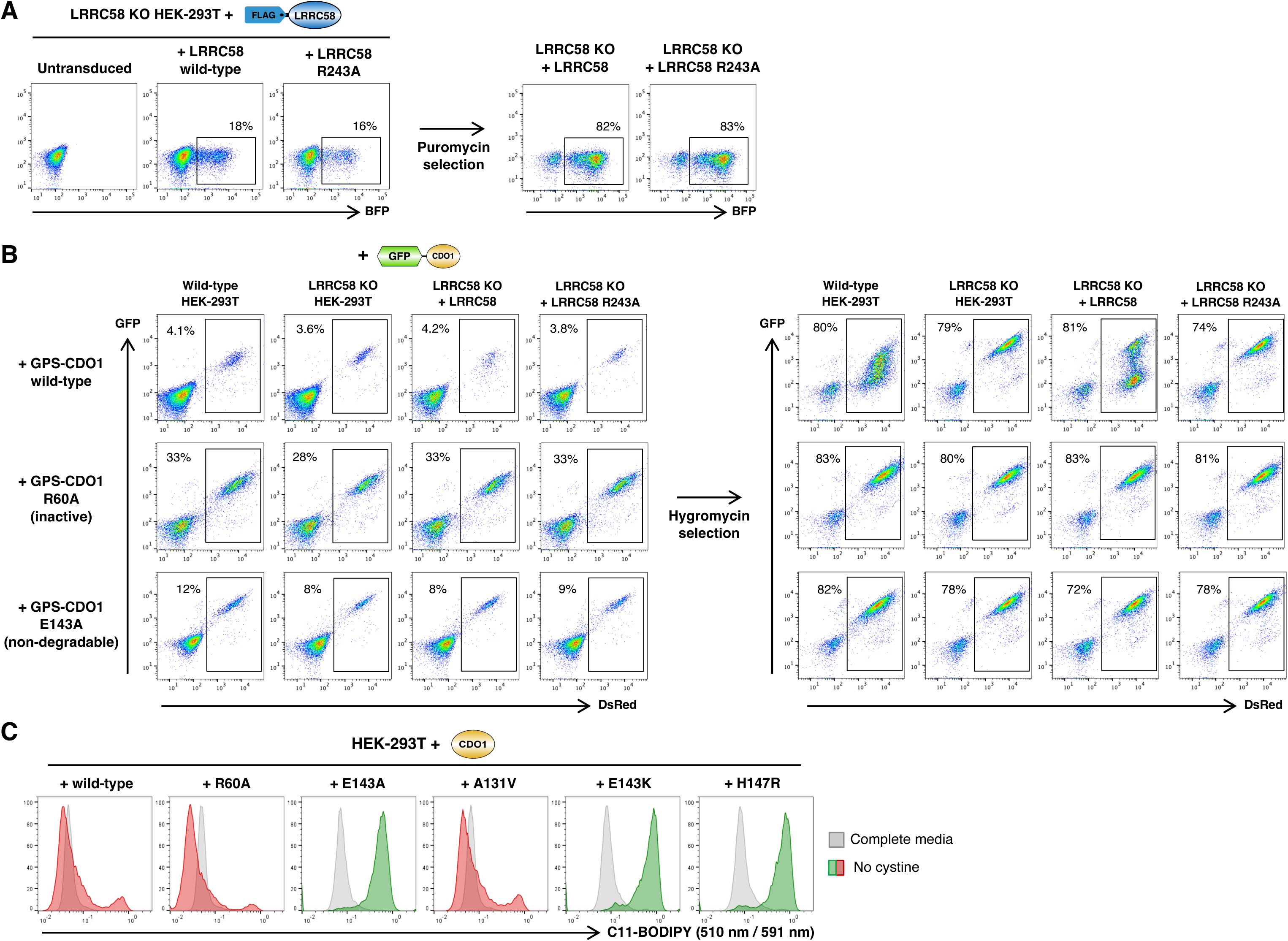
Manipulating the CDO1-LRRC58 axis in HEK-293T cells. **(A)** Starting with an LRRC58 knockout HEK-293T clone (thus expressing neither endogenous LRRC58 nor CDO1), the cells were transduced at single copy with lentiviral vectors encoding either wild-type LRRC58 or the R243A mutant of LRRC58 which is unable to bind CDO1. Untransduced cells were subsequently eliminated through puromycin selection. **(B)** The resulting cells, together with wild-type HEK-293T and the parental LRRC58 KO clone, were subsequently transduced at single copy with lentiviral vectors encoding either wild-type CDO1, R60A CDO1 which is catalytically-inactive, or E143A CDO1 which is unable to bind LRRC58 and purified via hygromycin selection. **(C)** CDO1 mutants refractory to LRRC58-mediated regulation induce lipid peroxidation upon cystine withdrawal. HEK-293T cells expressing the indicated CDO1 variants cultured in the presence and absence of cystine were stained with C11-BODIPY and analyzed by flow cytometry. The ratio of the green (510 nm) signal to the red (591 nm) signal is depicted.

**Supplementary Figure 7.**
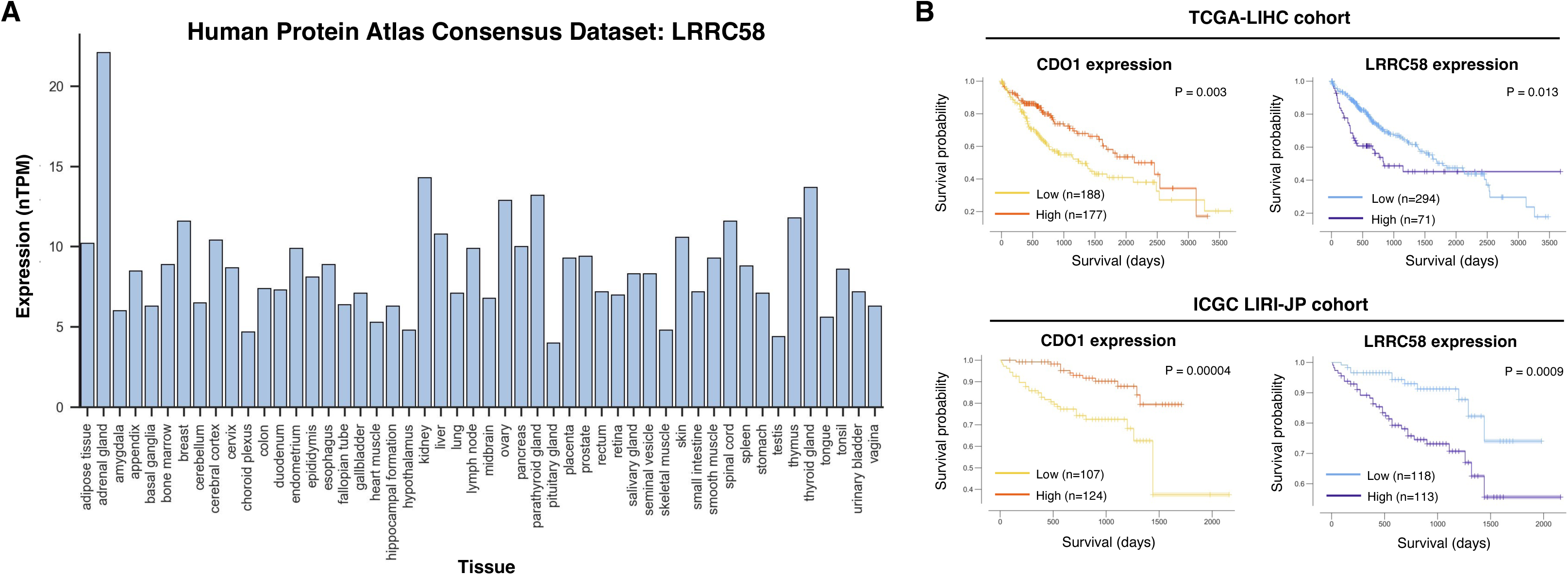
Expression pattern of LRRC58. **(A)** LRRC58 expression across human tissues as measured by RNA-seq, assessed using the consensus normalized expression (“nTPM”) values from the Human Protein Atlas. **(B)** Kaplan-Meier survival analysis of TCGA expression data revealing that high CDO1 expression and low LRRC58 expression is favorable in hepatocellular carcinoma.

## SUPPLEMENTARY TABLE LEGENDS

**Supplementary Table 1 | A stability profiling screen using the GPS-ORFeome library in Jurkat cells.** ORF-level data showing the corrected read counts across the six stability bins for each ORF detected in Jurkat. Analogous data generated previously in HEK-293T cells is derived from^25^, allowing the derivation of a ΔPSI metric reflecting the difference in stability between the two cell types. (PSI, protein stability index; 1 = maximally unstable, 6 = maximally stable).

**Supplementary Table 2 | A ubiquitin-focused CRISPR screen to identify genes required for the degradation of CDO1 upon cystine deprivation.** The output from the MAGeCK algorithm is shown. The top 50 genes are color-coded as follows: blue, Cul2 complex components; gray, proteasome subunits; red, p97 complex components; green, COP9 signalosome components.

**Supplementary Table 3 | Saturation mutagenesis stability profiling of CDO1 identifies residues required for instability in the absence of cysteine.** Raw data is provided for all mutants detected in the stability profiling screen. (PSI, protein stability index; 1 = maximally unstable, 4 = maximally stable).

**Supplementary Table 4 | TMT proteomics measuring protein abundance following withdrawal of cystine and methionine.** Raw data for all proteins quantified is shown.

**Supplementary Table 5 | A ubiquitin-focused CRISPR screen to identify genes required for the degradation of LRRC58 in complete media.** The output from the MAGeCK algorithm is shown. Cul2 complex components are indicated in blue.

**Supplementary Table 6 | Saturation mutagenesis stability profiling of LRRC58 identifies residues required for instability in complete media.** Raw data is provided for all mutants detected in the stability profiling screen. (PSI, protein stability index; 1 = maximally unstable, 3 = maximally stable).

**Supplementary Table 7 | Saturation mutagenesis activity profiling of LRRC58 identifies mutations which hyperactivate the degradation of CDO1 in complete media.** Raw data is provided for all mutants detected in the stability profiling screen. (PSI, protein stability index; 1 = maximally unstable, 3 = maximally stable).

## REFERENCES

1. González, A., and Hall, M.N. (2017). Nutrient sensing and TOR signaling in yeast and mammals. EMBO J 36, 397–408. 10.15252/EMBJ.201696010.

2. Efeyan, A., Comb, W.C., and Sabatini, D.M. (2015). Nutrient-sensing mechanisms and pathways. Nature 517, 302–310. 10.1038/NATURE14190.

3. Dong, J., Qiu, H., Garcia-Barrio, M., Anderson, J., and Hinnebusch, A.G. (2000). Uncharged tRNA activates GCN2 by displacing the protein kinase moiety from a bipartite tRNA-binding domain. Mol Cell 6, 269–279. 10.1016/S1097-2765(00)00028-9.

4. Masson, G.R. (2019). Towards a model of GCN2 activation. Biochem Soc Trans 47, 1481–1488. 10.1042/BST20190331.

5. Saxton, R.A., and Sabatini, D.M. (2017). mTOR Signaling in Growth, Metabolism, and Disease. Cell 168, 960–976. 10.1016/J.CELL.2017.02.004.

6. Wolfson, R.L., Chantranupong, L., Saxton, R.A., Shen, K., Scaria, S.M., Cantor, J.R., and Sabatini, D.M. (2016). Sestrin2 is a leucine sensor for the mTORC1 pathway. Science 351, 43–48. 10.1126/SCIENCE.AAB2674.

7. Saxton, R.A., Knockenhauer, K.E., Wolfson, R.L., Chantranupong, L., Pacold, M.E., Wang, T., Schwartz, T.U., and Sabatini, D.M. (2016). Structural basis for leucine sensing by the Sestrin2-mTORC1 pathway. Science (1979) 351, 53–58. 10.1126/science.aad2087.

8. Chantranupong, L., Scaria, S.M., Saxton, R.A., Gygi, M.P., Shen, K., Wyant, G.A., Wang, T., Harper, J.W., Gygi, S.P., and Sabatini, D.M. (2016). The CASTOR Proteins Are Arginine Sensors for the mTORC1 Pathway. Cell 165, 153–164. 10.1016/j.cell.2016.02.035.

9. Saxton, R.A., Chantranupong, L., Knockenhauer, K.E., Schwartz, T.U., and Sabatini, D.M. (2016). Mechanism of arginine sensing by CASTOR1 upstream of mTORC1. Nature 536, 229–233. 10.1038/NATURE19079.

10. Gu, X., Orozco, J.M., Saxton, R.A., Condon, K.J., Liu, G.Y., Krawczyk, P.A., Scaria, S.M., Wade Harper, J., Gygi, S.P., and Sabatini, D.M. (2017). SAMTOR is an S-adenosylmethionine sensor for the mTORC1 pathway. Science 358, 813–818. 10.1126/SCIENCE.AAO3265.

11. Tang, X., Zhang, Y., Wang, G., Zhang, C., Wang, F., Shi, J., Zhang, T., and Ding, J. (2022). Molecular mechanism of S-adenosylmethionine sensing by SAMTOR in mTORC1 signaling. Sci Adv 8. 10.1126/sciadv.abn3868.

12. Goul, C., Peruzzo, R., and Zoncu, R. (2023). The molecular basis of nutrient sensing and signalling by mTORC1 in metabolism regulation and disease. Nat Rev Mol Cell Biol 24, 857–875. 10.1038/S41580-023-00641-8.

13. Jin, J., Meng, T., Yu, Y., Wu, S., Jiao, C.C., Song, S., Li, Y.X., Zhang, Y., Zhao, Y.Y., Li, X., et al. (2025). Human HDAC6 senses valine abundancy to regulate DNA damage. Nature 637, 215–223. 10.1038/S41586-024-08248-5.

14. Chen, M.Y., Sun, C.Y., Zhao, R., Guan, X.L., Li, M.L., Zhang, F., Wan, Z.H., Feng, J.X., Yin, M., Lei, Q.Y., et al. (2025). BAG2 releases SAMD4B upon sensing of arginine deficiency to promote tumor cell survival. Mol Cell 85, 2581–2596.e6. 10.1016/j.molcel.2025.05.035.

15. Hershko, A., and Ciechanover, A. (1998). THE UBIQUITIN SYSTEM. Annu Rev Biochem 67, 425–479. 10.1146/annurev.biochem.67.1.425.

16. Zhang, Z., Mena, E.L., Timms, R.T., Koren, I., and Elledge, S.J. (2025). Degrons: defining the rules of protein degradation. Nat Rev Mol Cell Biol. 10.1038/s41580-025-00870-z.

17. Rape, M. (2018). Ubiquitylation at the crossroads of development and disease. Nat Rev Mol Cell Biol 19, 59–70. 10.1038/NRM.2017.83.

18. Gawden-Bone, C.M., Lehner, P.J., and Volkmar, N. (2023). As a matter of fat: Emerging roles of lipid-sensitive E3 ubiquitin ligases. Bioessays 45. 10.1002/BIES.202300139.

19. Menzies, S.A., Volkmar, N., van den Boomen, D.J.H., Timms, R.T., Dickson, A.S., Nathan, J.A., and Lehner, P.J. (2018). The sterol-responsive RNF145 E3 ubiquitin ligase mediates the degradation of HMG-CoA reductase together with gp78 and hrd1. Elife 7. 10.7554/eLife.40009.

20. Scott, N.A., Sharpe, L.J., and Brown, A.J. (2021). The E3 ubiquitin ligase MARCHF6 as a metabolic integrator in cholesterol synthesis and beyond. Biochim Biophys Acta Mol Cell Biol Lipids 1866. 10.1016/J.BBALIP.2020.158837.

21. Yen, H.C.S., Xu, Q., Chou, D.M., Zhao, Z., and Elledge, S.J. (2008). Global protein stability profiling in mammalian cells. Science (1979) *322*, 918–923. 10.1126/science.1160489.

22. Koren, I., Timms, R.T., Kula, T., Xu, Q., Li, M.Z., and Elledge, S.J. (2018). The Eukaryotic Proteome Is Shaped by E3 Ubiquitin Ligases Targeting C-Terminal Degrons. Cell 173, 1622–1635. 10.1016/j.cell.2018.04.028.

23. Stipanuk, M.H. (2020). Metabolism of sulfur-containing amino acids: How the body copes with excess methionine, cysteine, and sulfide. Journal of Nutrition 150, 2494S–2505S. 10.1093/jn/nxaa094.

24. Rowe, L., Mullegama, S. V., Lombardo, R., Barnes, C., Towner, S., Snyder, M.T., Heidlebaugh, A., Riordan, H., Begtrup, A., Crunk, A., et al. (2025). A proposed role for CDO1 in central nervous system development: Three children with rare missense variants and a neurological phenotype. HGG Adv, 100417. 10.1016/J.XHGG.2025.100417.

25. Timms, R.T., Mena, E.L., Leng, Y., Li, M.Z., Tchasovnikarova, I.A., Koren, I., and Elledge, S.J. (2023). Defining E3 ligase–substrate relationships through multiplex CRISPR screening. Nature Cell Biology 2023 25:10 25, 1535–1545. 10.1038/s41556-023-01229-2.

26. Paulsen, C.E., and Carroll, K.S. (2013). Cysteine-mediated redox signaling: chemistry, biology, and tools for discovery. Chem Rev 113, 4633–4679. 10.1021/CR300163E.

27. Poole, L.B. (2014). The Basics of Thiols and Cysteines in Redox Biology and Chemistry. Free Radic Biol Med 0, 148. 10.1016/J.FREERADBIOMED.2014.11.013.

28. Bonifácio, V.D.B., Pereira, S.A., Serpa, J., and Vicente, J.B. (2021). Cysteine metabolic circuitries: druggable targets in cancer. Br J Cancer 124, 862–879. 10.1038/S41416-020-01156-1.

29. Stipanuk, M.H., Hirschberger, L.L., Londono, M.P., Cresenzi, C.L., and Yu, A.F. (2004). The ubiquitin-proteasome system is responsible for cysteine-responsive regulation of cysteine dioxygenase concentration in liver. Am J Physiol Endocrinol Metab 286. 10.1152/AJPENDO.00336.2003.

30. Yoon, S.J., Combs, J.A., Falzone, A., Prieto-Farigua, N., Caldwell, S., Ackerman, H.D., Flores, E.R., and DeNicola, G.M. (2023). Comprehensive Metabolic Tracing Reveals the Origin and Catabolism of Cysteine in Mammalian Tissues and Tumors. Cancer Res 83, 1426. 10.1158/0008-5472.CAN-22-3000.

31. Gout, P.W., Buckley, A.R., Simms, C.R., and Bruchovsky, N. (2001). Sulfasalazine, a potent suppressor of lymphoma growth by inhibition of the x(c)-cystine transporter: a new action for an old drug. Leukemia 15, 1633–1640. 10.1038/SJ.LEU.2402238.

32. Yan, R., Xie, E., Li, Y., Li, J., Zhang, Y., Chi, X., Hu, X., Xu, L., Hou, T., Stockwell, B.R., et al. (2022). The structure of erastin-bound xCT-4F2hc complex reveals molecular mechanisms underlying erastin-induced ferroptosis. Cell Res 32, 687–690. 10.1038/S41422-022-00642-W.

33. Thoreen, C.C., Kang, S.A., Chang, J.W., Liu, Q., Zhang, J., Gao, Y., Reichling, L.J., Sim, T., Sabatini, D.M., and Gray, N.S. (2009). An ATP-competitive Mammalian Target of Rapamycin Inhibitor Reveals Rapamycin-resistant Functions of mTORC1. J Biol Chem 284, 8023. 10.1074/JBC.M900301200.

34. Dominy, J.E., Hwang, J., Guo, S., Hirschberger, L.L., Zhang, S., and Stipanuk, M.H. (2008). Synthesis of Amino Acid Cofactor in Cysteine Dioxygenase Is Regulated by Substrate and Represents a Novel Post-translational Regulation of Activity. J Biol Chem 283, 12188. 10.1074/JBC.M800044200.

35. Stipanuk, M.H., Londono, M., Lee, J.I., Hu, M., and Yu, A.F. (2002). Enzymes and metabolites of cysteine metabolism in nonhepatic tissues of rats show little response to changes in dietary protein or sulfur amino acid levels. J Nutr 132, 3369–3378. 10.1093/JN/132.11.3369.

36. Mahrour, N., Redwine, W.B., Florens, L., Swanson, S.K., Martin-Brown, S., Bradford, W.D., Staehling-Hampton, K., Washburn, M.P., Conaway, R.C., and Conaway, J.W. (2008). Characterization of Cullin-box sequences that direct recruitment of Cul2-Rbx1 and Cul5-Rbx2 modules to Elongin BC-based ubiquitin ligases. J Biol Chem 283, 8005–8013. 10.1074/JBC.M706987200.

37. Huttlin, E.L., Bruckner, R.J., Paulo, J.A., Cannon, J.R., Ting, L., Baltier, K., Colby, G., Gebreab, F., Gygi, M.P., Parzen, H., et al. (2017). Architecture of the human interactome defines protein communities and disease networks. Nature 545, 505–509. 10.1038/NATURE22366.

38. Jumper, J., Evans, R., Pritzel, A., Green, T., Figurnov, M., Ronneberger, O., Tunyasuvunakool, K., Bates, R., Žídek, A., Potapenko, A., et al. (2021). Highly accurate protein structure prediction with AlphaFold. Nature 2021 596:7873 596, 583–589. 10.1038/s41586-021-03819-2.

39. Zhang, Z., Sie, B., Chang, A., Leng, Y., Nardone, C., Timms, R.T., and Elledge, S.J. (2023). Elucidation of E3 ubiquitin ligase specificity through proteome-wide internal degron mapping. Mol Cell 83, 3377–3392.e6. 10.1016/J.MOLCEL.2023.08.022.

40. Ye, S., Wu, X., Wei, L., Tang, D., Sun, P., Bartlam, M., and Rao, Z. (2007). An Insight into the Mechanism of Human Cysteine Dioxygenase: KEY ROLES OF THE THIOETHER-BONDED TYROSINE-CYSTEINE COFACTOR. Journal of Biological Chemistry 282, 3391–3402. 10.1074/JBC.M609337200.

41. Poltorack, C.D., and Dixon, S.J. (2022). Understanding the role of cysteine in ferroptosis: progress & paradoxes. FEBS J 289, 374–385. 10.1111/FEBS.15842.

42. Dixon, S.J., Lemberg, K.M., Lamprecht, M.R., Skouta, R., Zaitsev, E.M., Gleason, C.E., Patel, D.N., Bauer, A.J., Cantley, A.M., Yang, W.S., et al. (2012). Ferroptosis: An iron-dependent form of nonapoptotic cell death. Cell 149, 1060–1072. 10.1016/j.cell.2012.03.042.

43. Drummen, G.P.C., Van Liebergen, L.C.M., Op den Kamp, J.A.F., and Post, J.A. (2002). C11-BODIPY(581/591), an oxidation-sensitive fluorescent lipid peroxidation probe: (micro)spectroscopic characterization and validation of methodology. Free Radic Biol Med 33, 473–490.

44. Hirschberger, L.L., Daval, S., Stover, P.J., and Stipanuk, M.H. (2001). Murine cysteine dioxygenase gene: structural organization, tissue-specific expression and promoter identification. Gene 277, 153–161. 10.1016/S0378-1119(01)00691-6.

45. De Bie, P., and Ciechanover, A. (2011). Ubiquitination of E3 ligases: self-regulation of the ubiquitin system via proteolytic and non-proteolytic mechanisms. Cell Death Differ 18, 1393. 10.1038/CDD.2011.16.

46. Sharpe, L.J., Howe, V., Scott, N.A., Luu, W., Phan, L., Berk, J.M., Hochstrasser, M., and Brown, A.J. (2018). Cholesterol increases protein levels of the E3 ligase MARCH6 and thereby stimulates protein degradation. J Biol Chem 294, 2436. 10.1074/JBC.RA118.005069.

47. Xiao, H., Ordonez, M., Fink, E.C., Covington, T.A., Woldemichael, H.B., Chen, J., Jain, M.S., Rohatgi, M.H., Wei, S.M., Burger, N., et al. (2025). Covariation MS uncovers a protein that controls cysteine catabolism. Nature 2025, 1–9. 10.1038/S41586-025-09535-5.

48. Combs, J.A., and Denicola, G.M. (2019). The Non-Essential Amino Acid Cysteine Becomes Essential for Tumor Proliferation and Survival. Cancers (Basel) 11. 10.3390/CANCERS11050678.

49. Paul, B.D., Sbodio, J.I., and Snyder, S.H. (2018). Cysteine metabolism in neuronal redox homeostasis. Trends Pharmacol Sci 39, 513. 10.1016/J.TIPS.2018.02.007.

50. Christensen, J.B., Donovan, A.P.A., Moradi, M., Vanacore, G., Helmy, M., Reid, A.J., Lee, J.T.H., Bayraktar, O.A., Brand, A.H., and Bayin, N.S. (2025). A conserved differentiation program facilitates inhibitory neuron production in the developing mouse and human cerebellum. bioRxiv, 2025.04.10.648162. 10.1101/2025.04.10.648162.

51. Sanjana, N.E., Shalem, O., and Zhang, F. (2014). Improved vectors and genome-wide libraries for CRISPR screening. Nat Methods 11, 783–784. 10.1038/nmeth.3047.

52. Li, W., Xu, H., Xiao, T., Cong, L., Love, M.I., Zhang, F., Irizarry, R.A., Liu, J.S., Brown, M., and Liu, X.S. (2014). MAGeCK enables robust identification of essential genes from genome-scale CRISPR/Cas9 knockout screens. Genome Biol 15, 554. 10.1186/s13059-014-0554-4.

53. Tzelepis, K., Koike-Yusa, H., De Braekeleer, E., Li, Y., Metzakopian, E., Dovey, O.M., Mupo, A., Grinkevich, V., Li, M., Mazan, M., et al. (2016). A CRISPR Dropout Screen Identifies Genetic Vulnerabilities and Therapeutic Targets in Acute Myeloid Leukemia. Cell Rep 17, 1193–1205. 10.1016/j.celrep.2016.09.079.

54. Clement, K., Rees, H., Canver, M.C., Gehrke, J.M., Farouni, R., Hsu, J.Y., Cole, M.A., Liu, D.R., Joung, J.K., Bauer, D.E., et al. (2019). CRISPResso2 provides accurate and rapid genome editing sequence analysis. Nat Biotechnol 37, 224–226. 10.1038/S41587-019-0032-3.

55. Ravenhill, B.J., Oliveira, M., Wood, G., Di, Y., Kite, J., Wang, X., Davies, C.T.R., Lu, Y., Antrobus, R., Elliott, G., et al. (2025). Spatial proteomics identifies a CRTC-dependent viral signaling pathway that stimulates production of interleukin-11. Cell Rep 44. 10.1016/J.CELREP.2025.115263,.

